# Identification of ligands binding to MB327-PAM-1, a binding pocket relevant for resensitization of nAChRs

**DOI:** 10.1101/2023.12.21.572862

**Authors:** Jesko Kaiser, Christoph G.W. Gertzen, Tamara Bernauer, Valentin Nitsche, Georg Höfner, Karin V. Niessen, Thomas Seeger, Franz F. Paintner, Klaus T. Wanner, Dirk Steinritz, Franz Worek, Holger Gohlke

## Abstract

Desensitization of nicotinic acetylcholine receptors (nAChRs) can be induced by overstimulation with acetylcholine (ACh) caused by an insufficient degradation of ACh after poisoning with organophosphorus compounds (OPCs). Currently, there is no generally applicable treatment for OPC poisoning that directly targets the desensitized nAChR. The bispyridinium compound MB327, an allosteric modulator of nAChR, has been shown to act as a resensitizer of nAChRs, indicating that drugs binding directly to nAChRs can have beneficial effects after OPC poisoning. However, MB327 also acts as an inhibitor of nAChRs at higher concentrations and can thus not be used for OPC poisoning treatment. Consequently, novel, more potent resensitizers are required. To successfully design novel ligands, the knowledge of the binding site is of utmost importance. Recently, we performed *in silico* studies to identify a new potential binding site of MB327, MB327-PAM-1, for which a more affine ligand, UNC0646, has been described. In this work, we performed ligand-based screening approaches to identify novel analogs of UNC0646 to help further understand the structure-affinity relationship of this compound class. Furthermore, we used structure-based screenings and identified compounds representing four new chemotypes binding to MB327-PAM-1. One of these compounds, cycloguanil, is the active metabolite of the antimalaria drug proguanil and shows a higher affinity towards MB327-PAM-1 than MB327. Furthermore, cycloguanil can reestablish the muscle force in soman-inhibited rat muscles. These results can act as a starting point to develop more potent resensitizers of nAChR and to close the gap in the treatment after OPC poisoning.

## Introduction

Chemical warfare agents remain a serious threat to the military and civilian population. Organophosphorus compounds (OPCs) are one class of chemical warfare agents and block acetylcholinesterase (AChE) covalently [1]. This inhibits the decomposition of acetylcholine causing inflated acetylcholine (ACh) concentrations in the synaptic gap and, thereby, an overstimulation of muscarinic (mAChR) and nicotinic (nAChR) acetylcholine receptors. The overstimulation leads to structural rearrangements of nAChR, resulting in a non-functional, desensitized state [1-3].

While the overstimulation of mAChRs can be treated with atropine, only antidotes (oximes) with insufficient efficiency are available to treat the overstimulation of nAChRs indirectly by reactivating AChE. However, these oximes are ineffective against several OPCs, in particular, when aging leads to altered OPC-enzyme complexes that are less susceptible to reactivation [4, 5]. Thus, novel antidotes are required to treat OPC poisonings.

The compound MB327 can reestablish the muscle force of OPC-poisoned muscles by interacting directly with nAChRs from several species via an allosteric modulation [6-10]. Furthermore, administration of MB327 can prolong the survival rates of guinea pigs after tabun poisoning compared to the oxime HI-6, both in combination with physostigmine and hyoscine [11]. While these results are promising, MB327 cannot be used in the treatment of OPC poisoning so far because the therapeutic range is too narrow. Restoration of the muscle force in a rat diaphragm myography assay by MB327 increases up to concentrations of 300 μM but decreases at higher concentrations [9, 10]. Similarly, MB327 initially shows positive allosteric effects on nicotine binding, which decrease at micromolar concentrations. Likewise, MB327 increases the binding of the orthosteric ligand epibatidine up to micromolar concentrations but at higher concentrations decreases it [7, 9]. These results indicate that MB327 transmits inhibitory effects on nAChR via binding to the orthosteric binding site, which has recently been experimentally validated for related bispyridinium compounds [12]. Additionally, we observed the binding of MB327 to the orthosteric binding site using free ligand diffusion molecular dynamics (MD) simulations [13]. Hence, novel compounds that are more affine and more selective to the allosteric binding site than MB327 need to be identified. A first step in this direction was recently done by us by identifying UNC0646 as an allosteric nAChR ligand with a higher affinity than MB327 (Sichler et al., submitted to Tox. Lett. on the 1^st^ of August 2023).

One way to identify novel binders and improve the affinity of known ligands is by using computer-aided drug design methods [14]. In ligand-based screening, one can search for analogs based on two-(topological) or three-(structural) dimensional ligand representations. In two-dimensional similarity screening, the importance of the three-dimensional conformation in the binding site is not taken into account. This can lead to unsatisfactory results, especially for highly flexible ligands. Thus, three-dimensional ligand screening approaches can be more effective in identifying binders that can fit into the binding site and have been used for identifying novel binders successfully in the past [15-20]. However, ligand-based screening approaches neglect explicit knowledge of the receptor structure. Additionally, ligand-based screening may only identify hits with more similar chemical scaffolds compared to the query. Thus, structure-based screening is a popular approach to identifying novel ligands for biological targets and can help to identify novel chemical scaffolds [21-28]. Recently, we identified a novel allosteric binding site of MB327 (MB327-PAM-1) and described a potential binding mode of UNC0646 in MB327-PAM-1 [13, 29], providing necessary input for three-dimensional ligand-based screening and structure-based screening.

Here, we used this information to perform all three described screening approaches to increase the chance of success and validated hits by a mass spectrometry-based affinity determination (MS Binding Assay). We identified novel substituents (**1a-k, 2b-e, 2g**) at the UNC0646 quinazoline scaffold that lead to higher affinity and ligands with novel chemotypes [PTMD99-0001C (**3**), PTMD99-0016C (**4**), PTMD99-0026C (**5**), and cycloguanil (**6**)] binding to MB327-PAM-1.

## Materials and Methods

### Two-dimensional similarity search

The MolPort database was searched for compounds similar to UNC0646 using a two-dimensional search as implemented on the MolPort website (https://molport.com, accessed on October 21^st^, 2020). All compounds identified by the similarity search using default parameters were selected. From 22 compounds identified, twelve compounds were ordered for testing in this study.

### Homology modeling of nAChR

The homology modeling of the human and *Torpedo* (recently reclassified as *Tetronarce*) nAChR was performed as previously described [13]. In short, homology models were generated using MODELLER version 9.19 [30] with the PDB structures 6PV8 [31], 5KXI [32], 2WN9 [33], and 6CNK [34] as templates. Water molecules and molecules from the crystallization buffer were removed. Amino acids not resolved in the templates were not included in the models; these amino acids are located within the intracellular loop, the *N-*and *C-*terminal region and, hence, not in the region that forms the MB327-PAM-1 pocket. The final models were selected based on the DOPE potential [35], TopScore [36], and visual inspection and subsequently protonated using PROPKA, v3.4.0 [37, 38] as implemented in Schrödinger Maestro, v2021-1 [39] at pH 7.4. The termini were capped with *N*-methyl amide (NME) and acetyl (ACE) groups, respectively.

### Ligand-based screening with subsequent template-based docking

Based on our proposed binding mode of UNC0646 [29], we used the binding mode of PTMD01-0004 (**2a**) [Bernauer *et al.*, in preparation], a structurally reduced analog of UNC0646 that lacks the quinazoline substituent in 2-position, as input for ligand-based screening. We created a database from feasible organic reactions of building blocks of PTMD01-0004 (**2a**) with MolPort (https://molport.com) building blocks as implemented in the PINGUI webserver (https://scubidoo.pharmazie.uni-marburg.de/pingui/, accessed on May 19^th^, 2021) (SI Figure S1) [40], resulting in 69,223 in principle synthesizable compounds. We protonated the constituents of the database using OpenEye FixpKa, v2.1.1.0 [41] and filtered the database to only use compounds that are at least double positively charged as PTMD01-0004 (**2a**), resulting in 14,396 compounds and generated conformers using OpenEye OMEGA, v4.1.1.0 [42, 43] with default parameters except setting the *strictstereo* parameter to false. Initially, we used OpenEye vROCS, v3.4.1.0 [44, 45] (SI Figure S2) to filter the database by applying default parameters and the TanimotoCombo score. The best 1000 hits from the vROCS screening were further investigated using CCG MOE, v2020 [46] using a template-based docking with an upstream pharmacophore filter (SI Figure S3). The hits were analyzed based on visual inspection, and four compounds were selected for synthesizing.

### Synthesis

All target compounds synthesized in the context of this study were cataloged with a PTMD number (<u>P</u>harmacy and <u>T</u>oxicology <u>M</u>unich and <u>D</u>üsseldorf). All chemicals were used as purchased from commercial sources. Solvents used for purification were distilled before use. 5-Pyrrolidin-1-ylpentan-1-amine, which could only be purchased as the respective hydrochloride salt was converted into the free base before use [47]. Anhydrous reactions were carried out under an argon atmosphere in vacuum-dried glassware. For microwave reactions, a Discover SP microwave system by CEM GmbH was used. TLC-Analysis was performed on plates purchased from Merck (silica gel 60F_254_ on aluminum sheet). Flash chromatography was carried out using silica gel 60 (40-63 mm mesh size) as stationary phase, purchased from Merck. All synthesized compounds were dried under a high vacuum. ^1^H and ^13^C NMR spectra were recorded with a Bruker BioSpin Avance III HD 400 and 500 MHz at 25 °C. For data processing, MestReNova (Version 14.1.0) from Mestrelab Research S.L. 2019, and for calibration, the solvent signal (CD_2_Cl_2_, CD_3_OD or DMSO-d_6_) was used. The purity of the test compounds was ≥ 95%, determined by means of quantitative NMR using TraceCERT® ethyl 4-(dimethylamino)benzoate from Merck as an internal calibrant [48, 49]. High-resolution mass spectra were recorded on a Finnigan MAT 95 (EI) or a Finnigan LTQ FT (ESI). Melting points were determined with a Büchi 510 melting point instrument and are uncorrected. For IR spectroscopy, an FT-IR Spectrometer 1600 from PerkinElmer was used. The analytical data of the synthesized compounds described below, obtained using the described methods, can be found in the Supporting Information.

Synthesis of 2,4-dichloro-6-methoxy-7-[3-(piperidin-1-yl)propoxy]quinazoline (**7**) recently reported by Vital *et al.* [50] and of *N*-(1-cyclohexylpiperidin-4-yl)-6-methoxy-7-[3-(piperidin-1-yl)propoxy]-quinazolin-4-amine (**2a**) was accomplished according to Bernauer *et al.* [Bernauer *et al.*, in preparation].

#### General Procedure: Synthesis of quinazolin-4-amines (GP)

A solution of the respective 4-chloroquinazoline (**8**) or (**9**) (1.0 equiv), the corresponding amine (2.0 equiv - 10 equiv) and *N*,*N*-diisopropylethylamine (DIEA) (3.0 equiv) in *i*-PrOH (5 mL/mmol) was stirred at 160 °C for 15 to 60 min under microwave irradiation (200 W). The reaction mixture was concentrated in vacuo and the crude product was purified by flash chromatography [5% to 15% 3 M NH_3_ (in MeOH) in CH_2_Cl_2_ or 10% MeOH in CH_2_Cl_2_ + 0.5% DMEA].

#### 6-Methoxy-2-(piperidin-1-yl)-7-[3-(piperidin-1-yl)propoxy]-*N*-[5-(pyrrolidin-1-yl)pentyl]quinazolin-4-amine (1k)

According to GP with **8** (126 mg, 0.300 mmol, 1.0 equiv), 5-(pyrrolidin-1-yl)pentan-1-amine (93.8 mg, 0.600 mmol, 2.0 equiv) and DIEA (160 µL, 119 mg, 0.900 mmol, 3.0 equiv) in *i-*PrOH (1.5 mL) for 15 min. **1k** (117 mg, 72% yield, 96% purity) was isolated by flash chromatography [7% to 15% 3 M NH_3_ (in MeOH) in CH_2_Cl_2_] as a colorless solid.

#### *N*^1^-Cyclohexyl-*N*^2^-{6-methoxy-7-[3-(piperidin-1-yl)propoxy]quinazolin-4-yl}-*N*^1^-methylethane-1,2-diamine (2b)

According to GP with **9** (101 mg, 0.300 mmol, 1.0 equiv), *N*^1^-cyclohexyl-*N*^1^-methylethane-1,2-diamine (104 µL, 93.8 mg, 0.600 mmol, 2.0 equiv) and DIEA (160 µL, 119 mg, 0.900 mmol, 3.0 equiv) in *i-*PrOH (1.5 mL) for 1 h. **2b** (127 mg, 93%) was isolated by flash chromatography [10% 3 M NH_3_ (in MeOH) in CH_2_Cl_2_] as a yellow oil (96% purity).

#### 6-Methoxy-*N*-[3-(4-methylpiperazin-1-yl)butyl]-7-[3-(piperidin-1-yl)propoxy]quinazolin-4-amine (2c)

According to GP with **9** (101 mg, 0.300 mmol, 1.0 equiv), 3-(4-methylpiperazin-1-yl)butan-1-amine (114 µL, 108 mg, 0.600 mmol, 2.0 equiv) and DIEA (160 µL, 119 mg, 0.900 mmol, 3.0 equiv) in *i*-PrOH (1.5 mL) for 1 h. **2c** (110 mg, 78% yield, 97% purity) was isolated by flash chromatography [10% 3 M NH_3_ (in MeOH) in CH_2_Cl_2_] as a pale yellow solid.

#### 6-Methoxy-7-[3-(piperidin-1-yl)propoxy]-4-[4-(pyrrolidin-1-yl)piperidin-1-yl]quinazoline (2d)

According to GP with **9** (101 mg, 0.300 mmol, 1.0 equiv), 4-(pyrrolidin-1-yl)piperidine (97.4 mg, 0.600 mmol, 2.0 equiv) and DIEA (160 µL, 119 mg, 0.900 mmol, 3.0 equiv) in *i-*PrOH (1.5 mL) for 1 h. **2d** (122 mg, 89%) was isolated by flash chromatography [10% 3 M NH_3_ (in MeOH) in CH_2_Cl_2_] as a yellow oil (97% purity).

#### *N*-[1-(Azepan-1-yl)-2-methylpropan-2-yl]-6-methoxy-7-[3-(piperidin-1-yl)propoxy]quinazolin-4-amine (2e)

According to GP with **9** (134 mg, 0.400 mmol, 1.0 equiv), 1-(azepan-1-yl)-2-methylpropan-2-amine (717 mg, 4.00 mmol, 10 equiv) and DIEA (213 µL, 158 mg, 1.20 mmol, 3.0 equiv) in *i-*PrOH (2.0 mL) for 1 h. **2e** (46.4 mg, 25% yield, 97% purity) was isolated by flash chromatography [1. 10% 7 M NH_3_ (in MeOH) in CH_2_Cl_2_, 2. 10% MeOH in CH_2_Cl_2_ + 0.5% DMEA] as a yellow oil. In addition, a further product fraction of lower purity was obtained (21% yield, 72% purity).

#### *N*-(1-Propan-2-ylpiperidin-4-yl)-6-methoxy-7-[3-(piperidin-1-yl)propoxy]quinazolin-4-amine (2f)

According to GP with **9** (33.6 mg, 0.100 mmol, 1.0 equiv), 1-propan-2-ylpiperidin-4-amine (31.6 µL, 28.4 mg, 0.200 mmol, 2.0 equiv) and DIEA (53.3 µL, 39.6 mg, 0.300 mmol, 3.0 equiv) in *i-*PrOH (0.5 mL) for 15 min. **2g** (41.2 mg, 93%) was isolated by flash chromatography [5% to 10% 3 M NH_3_ (in MeOH) in CH_2_Cl_2_] as a colorless solid (96% purity).

#### *N*-(1-Propan-2-ylpiperidin-4-yl)-6-methoxy-7-[3-(piperidin-1-ylmethyl)pyrrolidin-1-yl]quinazolin-4-amine (2g)

A mixture of **10** (111 mg, 0.350 mmol, 1.0 equiv), 1-(pyrrolidin-3-ylmethyl)piperidine (310 mg, 1.75 mmol, 5.0 equiv) and potassium carbonate (53.2 mg, 0.385 mmol, 1.1 equiv) in *N*-methyl-2-pyrrolidone (NMP) (455 µL) was stirred at 135 °C for 20 h. The reaction mixture was concentrated and purified by flash chromatography [5% to 20% 3 M NH_3_ (in MeOH) in CH_2_Cl_2_] to afford **2f** (147 mg, 90%) as a pale yellow solid (99% purity).

#### 4-Chloro-6-methoxy-2-(piperidin-1-yl)-7-[3-(piperidin-1-yl)propoxy]quinazoline (8)

A solution of 7 (370 mg, 1.00 mmol, 1.0 equiv) and 1-methylpiperidine (244 µL, 200 mg, 2.00 mmol, 2.0 equiv) in 1,4-dioxane (2.5 mL) was stirred at 150 °C for 1 h under microwave irradiation (300 W). **8** was isolated by flash chromatography (10% to 20% MeOH in CH_2_Cl_2_) as a yellow solid (327 mg, 78% yield, 95% purity).

#### 4-Chloro-6-methoxy-7-[3-(piperidin-1-yl)propoxy]quinazoline (9) [51, 52]

To a slurry of **11** (22.2 mg, 0.100 mmol, 1.0 equiv), 3-piperidin-1-ylpropan-1-ol (19.9 µL, 18.8 mg, 0.125 mmol, 1.25 equiv), PPh_3_ (34.4 mg, 0.130 mmol, 1.3 equiv) and dry THF (1.0 mL) was added di-*tert*-butyl azodicarboxylate (DBAD) (30.5 mg, 0.130 mmol, 1.3 equiv) in portions at 0 °C. The resulting solution was stirred overnight at rt and concentrated under reduced pressure. Purification by flash chromatography [5% 3 M NH_3_ (in MeOH) in CH_2_Cl_2_] afforded **9** as a pale yellow solid (34.5 mg, > 99% yield, 96% purity).

#### 7-Fluoro-N-(1-propan-2-ylpiperidin-4-yl)-6-methoxyquinazolin-4-amine (10)

A mixture of 12 (20.2 mg, 0.100 mmol, 1.0 equiv), PyBOP (69.0 mg, 0.130 mmol, 1.5 equiv), diazabicycloundecene (DBU) (22.9 µL, 23.3 mg, 0.150 mmol, 1.5 equiv) and 1-propan-2-ylpiperidin-4-amine (23.7 µL, 21.3 mg, 0.150 mmol, 1.5 equiv) in acetonitrile (0.5 mL) was stirred at rt for 1 h. The mixture was concentrated in vacuo. **10** (29.3 mg, 92%) was isolated after flash chromatography [5% 3 M NH_3_ (in MeOH) in CH_2_Cl_2_] as a colorless solid (97% purity).

### Structure-based screening

The SMILES codes of in-stock compounds in the lead-like (3,434,621 compounds) and the double-protonated (129,606 compounds) subsets were downloaded from the ZINC20 database [53]. All compounds were protonated using OpenEye FixpKa, v2.1.1.0, and conformers were generated using OpenEye OMEGA, v4.1.0.0 [42] with default parameters except setting the *strictstereo* parameter to false [42, 43]. The human and *Torpedo* homology models of nAChR were prepared for docking using OpenEye MakeReceptor, v4.0.0.0, and all compounds were docked using OpenEye FRED, v4.0.0.0 [54-56], writing out a maximum of one pose per compound. The best 1000 hits in each binding pocket were visually inspected, and 30 compounds were ordered for testing.

### Commercially obtained compounds

The 42 commercially obtainable compounds were purchased from several suppliers with purities of at least 85%. A corresponding, detailed list can be found in the Supporting Information (SI Table S1). For affinity testing in MS Binding Assays, the compounds were applied as described in SI Table S1.

### Molecular dynamics simulations of cycloguanil bound to nAChR

The receptor with the docked ligand was embedded in a membrane of 1-palmitoyl-2-oleoyl-*sn*-glycero-3-phosphocholine (POPC) lipids and solvated using Packmol-Memgen [57] in a rectangular box of TIP3P water [58]. KCl was added at a concentration of 150 mM and the system was neutralized using Cl^-^ ions. The edge of the box was set to be at least 12 Å away from receptor atoms. The AMBER22 package of molecular simulations software [59, 60] was used in combination with the *ff14SB* force field for the protein [61], the Lipid21 force field for the lipids [62], and the Joung and Cheatham parameters for monovalent ions [63]. Ligand charges were calculated according to the RESP procedure [64] with default parameters as implemented in antechamber [65] using electrostatic potentials generated by Gaussian16 [66] at the HF-6-31G* level of theory; ligand force field parameters were derived from the gaff2 force field. Since cycloguanil (**6**) should carry one positive charge (p*K*_a_ = 11.4 [67]) on the nitrogen atoms in the 1,6-dihydro-1,3,5-triazine-2,4-diamine ring system, *N*-3 was protonated.

Molecular dynamics (MD) simulations were performed as described earlier [13] using AMBER22. In short, a combination of steepest descent and conjugate gradient minimization was performed while lowering positional restraints with force constants from 25 kcal mol^-1^ Å^-2^ to zero. Stepwise heating to 300 K and a subsequent reduction of harmonic restraints from 25 kcal mol^-1^ Å^-2^ to zero followed.

Subsequently, 10 replicas of 1 μs length each of unbiased MD simulations were performed; for temperature control, the Langevin dynamics were applied with a collision frequency of 2 ps^-1^, and the Berendsen barostat with semi-isotropic pressure adaption was used.

Based on the four replicas in which cycloguanil (**6**) remained within the binding site, we computed representative binding structures of cycloguanil (**6**) using the *k*-means clustering algorithm, as implemented in CPPTRAJ [68]. We then restarted simulations from the representative binding mode. Because the hydrogen bonds to E62_γ_ and E200_γ_ were highly conserved among all clusters except the largest one (containing 18.3% of all frames), we decided to restart simulations from the second largest cluster (containing 14.1% of all frames) (SI Figure S4). Therefore, we started directly with the production run with similar settings as for the docked structure. The first 10 ns of each replica were removed for further analysis. All simulations were analyzed using CPPTRAJ [68].

### Prediction of physicochemical and toxicological properties

Using OpenEye OMEGA, version 4.1.1.1 [43, 69], three-dimensional conformations of the compounds based on the SMILES code were generated using the default setting with the exception that only one conformation was generated for each compound. Pharmacokinetic properties and hERG inhibition were predicted using Schrödinger QikProp, version 2022-2 [70].

For PAINS filtering, the PAINS-remover webserver, v0.99 (https://www.cbligand.org/PAINS/) [71], was used. For the prediction of further toxicological properties, we used NEXUS Derek, v6.0.1 [72].

### Image generation

Images of nAChR were created using UCSF Chimera [73].

### Rat diaphragm myography

All procedures using animals followed animal care regulations and were approved by the responsible ethics committee. Preparation of rat diaphragm hemispheres and experimental protocol of myography were performed as described before [9]. In short, for all procedures (including wash-out steps, preparation of soman and bispyridinium compound solutions) aerated Tyrode solution (125 mM NaCl, 24 mM NaHCO_3_, 5.4 mM KCl, 1 mM MgCl_2_, 1.8 mM CaCl_2_, 10 mM glucose, 95 % O_2_, 5% CO_2_; pH 7.4; 25 ± 0.5 ^◦^C) was used. After the recording of control muscle force, the muscle preparations were incubated in the Tyrode solution, containing 3 μM soman. Following a 20 min wash-out period, the test compound cycloguanil (Merck KGaA) was added in ascending concentrations (1 μM, 10 μM, 30 μM, 70 μM, 100 μM, 150 μM, 200 μM, 300 μM, 500 μM, 1000 μM). In each preparation, four concentrations were measured to avoid the fatigue effects of muscle force generation. The incubation time was 20 min for each concentration. The electric field stimulation was performed with 10 μs pulse width and 0.2 A amplitudes. The titanic trains of 20 Hz, 50 Hz, 100 Hz were applied for 1 s and in 10 min intervals. Measurements on non-toxic muscle were carried out according to the same scheme. Instead of soman, pure Tyrode was incubated. Muscle force was calculated as a time-force integral (area under the curve, AUC) and constrained to values obtained for maximal force generation (muscle force in the presence of Tyrode solution without any additives; 100 %).

### UNC0642 MS Binding Assays

Competitive MS Binding Assays were performed as described previously [29]. In short, the reporter ligand (UNC0642, 1 µM) and the corresponding test compound (varying concentrations) were incubated with aliquots of a membrane preparation from *Torpedo californica* electroplaque (approx. 75 µg protein per sample) in incubation buffer (120 mM NaCl, 5 mM KCl, 8.05 mM Na_2_HPO_4_ and 1.95 mM NaH_2_PO_4_, pH 7.4). Samples were generated in triplicates. After separating the protein-bound from non-bound reporter ligand by centrifugation, the protein-bound portion of UNC0642 was liberated and finally quantified via LC-ESI-MS/MS. Total binding of UNC0642 was normalized to 100% (i.e. reporter ligand binding in the absence of test compound) and 0% (i.e. non-specific reporter ligand binding, determined by the presence of 100 µM UNC0646 instead of test compound). Applying the non-linear regression function “One site – fit Ki” yielded competition curves, which then revealed IC_50_ and *K*_i_ values, respectively (Prism software v. 6.07, GraphPad software, La Jolla, CA, USA). Top and bottom levels were fixed at 100% and 0%, respectively, for that purpose. *K*_i_ values are given as mean p*K*_i_ values from three experiments ± SEM, if not stated otherwise. Next to the full-scale competition experiments, also competition experiments with only a single concentration of test compound (i.e. 10 µM) have been performed in this study. Compared to full-scale competition experiments, the obtained data was not analyzed via non-linear regression in this case and only normalized as described above to reveal the remaining reporter ligand binding as an indicator for the affinity of the respective test compound. If not stated otherwise, the remaining reporter ligand binding is given as the mean of triplicates ± SD.

## Results and Discussion

### Screening strategy

We used different strategies to identify novel binders of MB327-PAM-1. First, we performed ligand-based screening to identify analogs of UNC0646 using a two-dimensional similarity search as implemented on the MolPort website (https://molport.com) to identify compounds with high similarity to UNC0646 (Figure 1, blue scheme). Furthermore, we used PTMD01-0004 (**2a**), an analog with no substituent in the 2-position of the quinazoline ring, to perform ligand-based screening using its three-dimensional binding mode as a query using OpenEye vROCS [44, 45], followed by a pharmacophore-based docking of the best hits using CCG MOE [46] (Figure 1, yellow scheme). Second, to reveal new binders with novel chemical scaffolds, we performed structure-based virtual screenings (Figure 1, green scheme). There, we first docked a ZINC20 [53] lead-like library (3,434,621 compounds) into the human muscle-type nAChR using OpenEye FRED [54-56]. The lead-like library only includes compounds of a molecular weight between 250-350 g mol^-1^. However, because a higher affinity of known ligands binding to MB327-PAM-1 generally correlates with a larger size of the ligands and most previously described binders feature at least two positive charges [29, 74, 75] (Sichler *et al.,* submitted to Tox. Lett. on the 1^st^ of August 2023), we performed an additional screening using a ZINC20 library [53] of in-stock compounds bearing at least two positive charges (129,606 compounds). To exploit that the amino acids interacting with MB327 and UNC0646 are highly conserved among the human muscle-type and *Torpedo* nAChR [13, 29] but that the conformations of the sidechains nevertheless vary, we now used the *Torpedo* nAChR for docking to increase the search space. Also, the *Torpedo* nAChR is used in our MS Binding Assays.

**Figure 1:**
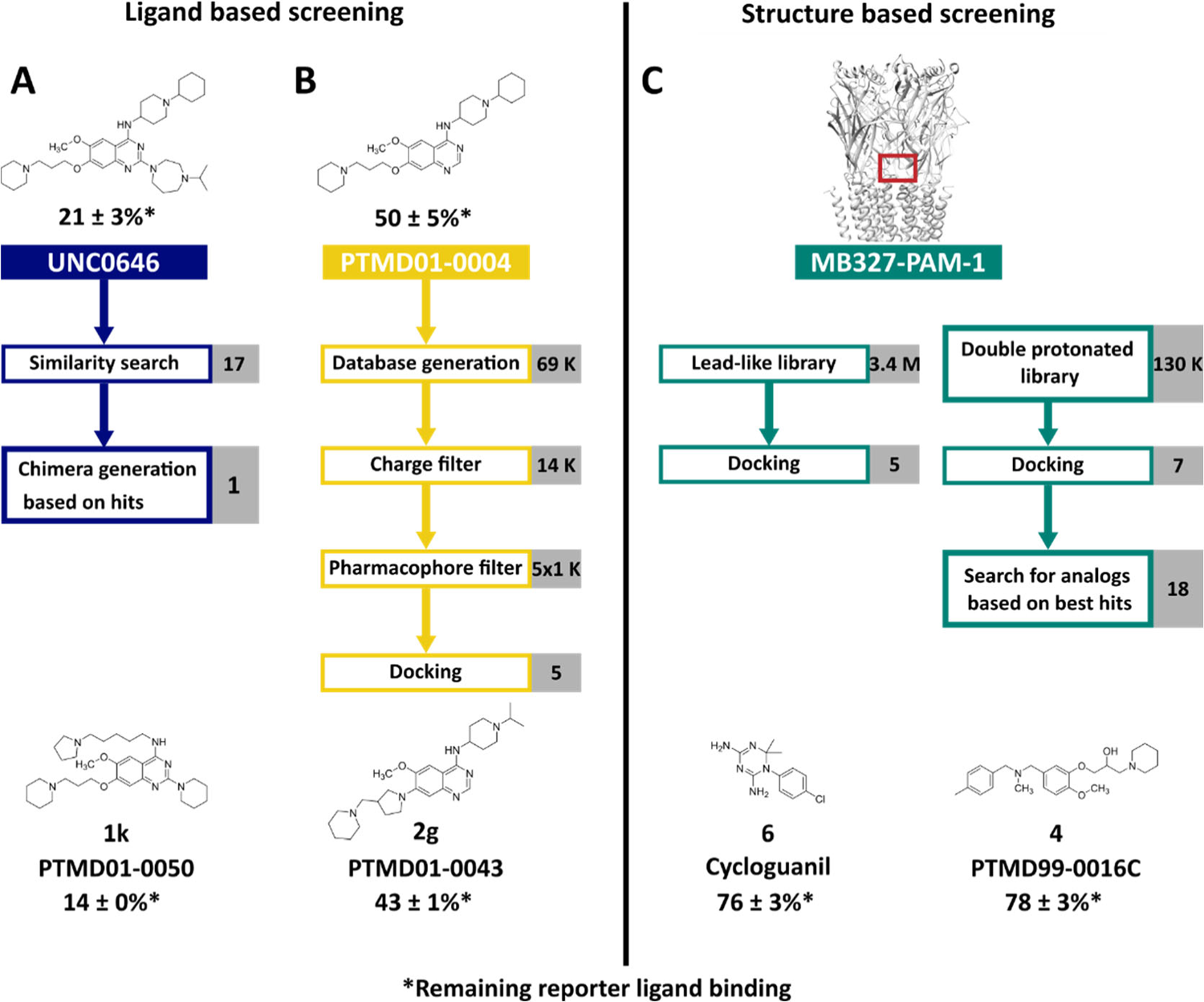
Screening strategies to identify novel binders of MB327-PAM-1. **A)** A two-dimensional similarity search using UNC0646 as a template was performed using MolPort (https://www.molport.com/). Based on the hits, one novel chimera compound was designed. **B)** Based on PTMD01-0004 (**2a**), a UNC0646-analog lacking the side chain in the 2-position, a database was generated based on feasible organic reactions (SI Figure S1). After applying a charge filter and a pharmacophore filter, docking experiments led to five novel analogs of UNC0646. **C)** Based on structure-based screening experiments in the human muscle type and *Torpedo* nAChR, 5 respectively 7 compounds with novel chemotypes were ordered for affinity characterization in MS Binding Assays. After the first experimental results, 18 additional compounds based on three chemical scaffolds were selected from the initial screening and ordered. This resulted in the identification of cycloguanil (**6**) and PTMD99-0016C (**4**). For each screening strategy, the best hits are shown. Percentage values indicate the remaining reporter ligand binding in the presence of test compounds (at 10 μM concentration) as compared to 100% reporter ligand binding in the absence of a competitor using the reporter ligand UNC0642 in MS Binding Assays (1 µM UNC0642) (mean ± SD, *n* = 3).

### Two-dimensional similarity search yields affine UNC0646 analogs with small substituents in 4-position

Based on a two-dimensional screening of the MolPort library using UNC0646 as a template (Figure 1, blue scheme), 12 compounds were tested in our MS Binding Assay for MB327-PAM-1 [29]. Of these, 10 compounds displaced the reporter ligand UNC0642 from the binding site at 10 μM (**1a**-**1j**; remaining reporter ligand binding at most 90 ± 7%, *n* = 3; Table 1, SI Table S2). The best result was obtained for PTMD01-0019C (**1a)**. In contrast to previously described UNC0646 analogs (Sichler et al., submitted to Tox. Lett. on the 1^st^ of August 2023) [29], this is the only compound that does not have a side chain in 7-position with an aliphatic amino group, although mainly acidic amino acid side chains are available for ligand binding in MB327-PAM-1 [13, 29]. However, a study with related 4-amino-2-(*N,N*-diethylamino)quinazoline derivatives revealed that the two amino substituents impact the p*K*_a_ values of quinazolines resulting in p*K*_a_ values of up to 8.31 [76] compared to 3.51 of the unsubstituted quinazoline [77]. Thus, PTMD01-0019C (**1a**) can still be protonated under experimental and physiological conditions, similar to all previously described binders in MB327-PAM-1 [13, 29, 74, 75]. Furthermore, 4-aminopyridine has a p*K*_a_ of 9.17, indicating that even the residue in the 2-position of PTMD01-0019C (**1a**) may be protonated [78]. These results are in line with suggestions that a positive charge is crucial for binding in MB327-PAM-1 but also indicate that the location of the positive charge is less important, which can be explained by the many acidic amino acids in MB327-PAM-1 [13, 29]. To verify the results obtained from the competition experiments by applying a single concentration, we performed full-scale competition studies for the best-binding compound PTMD01-0019C (**1a**), resulting in a p*K*_i_ of 5.19 ± 0.05 (SI Figure S5).

**Table 1:**
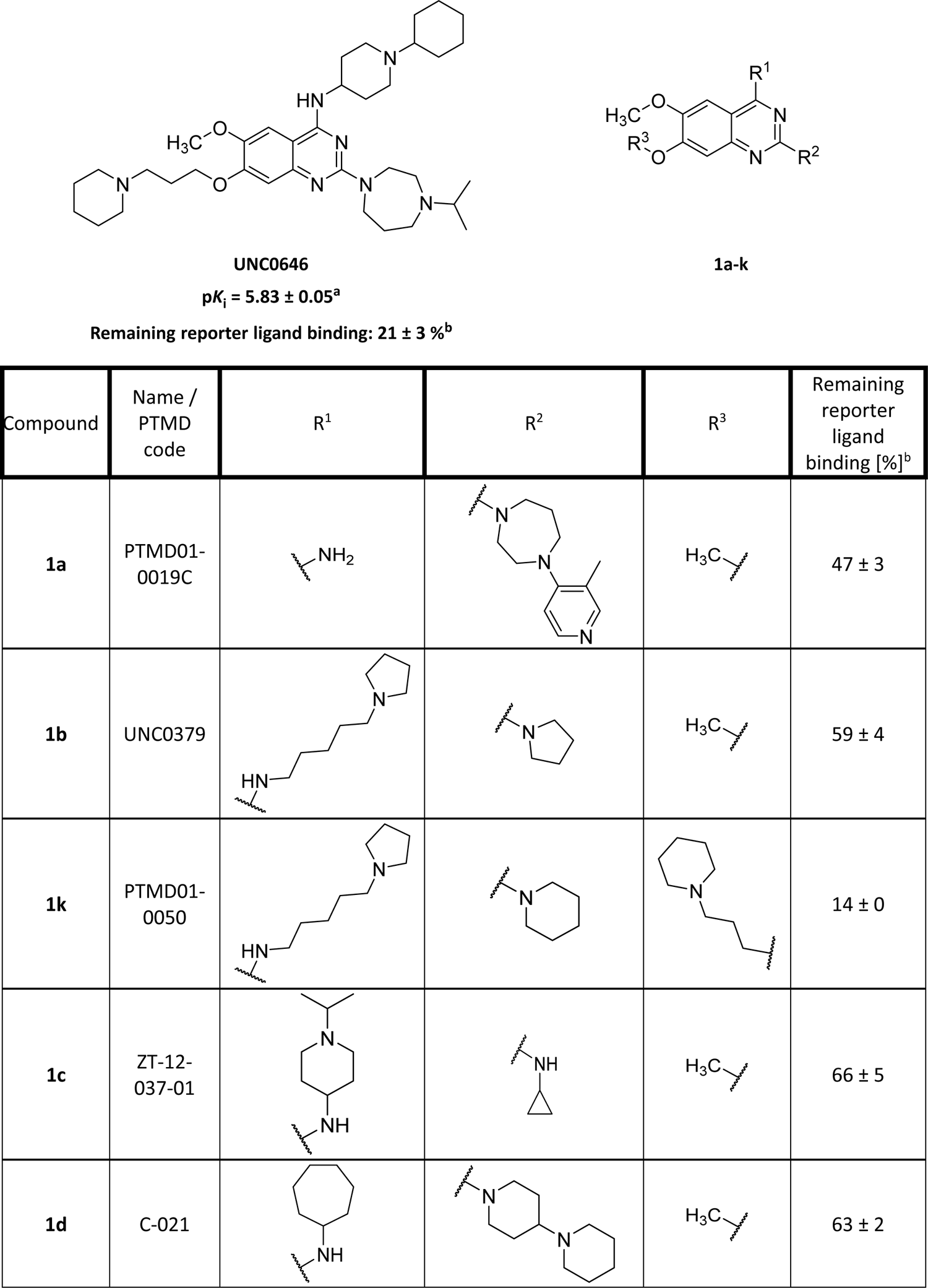
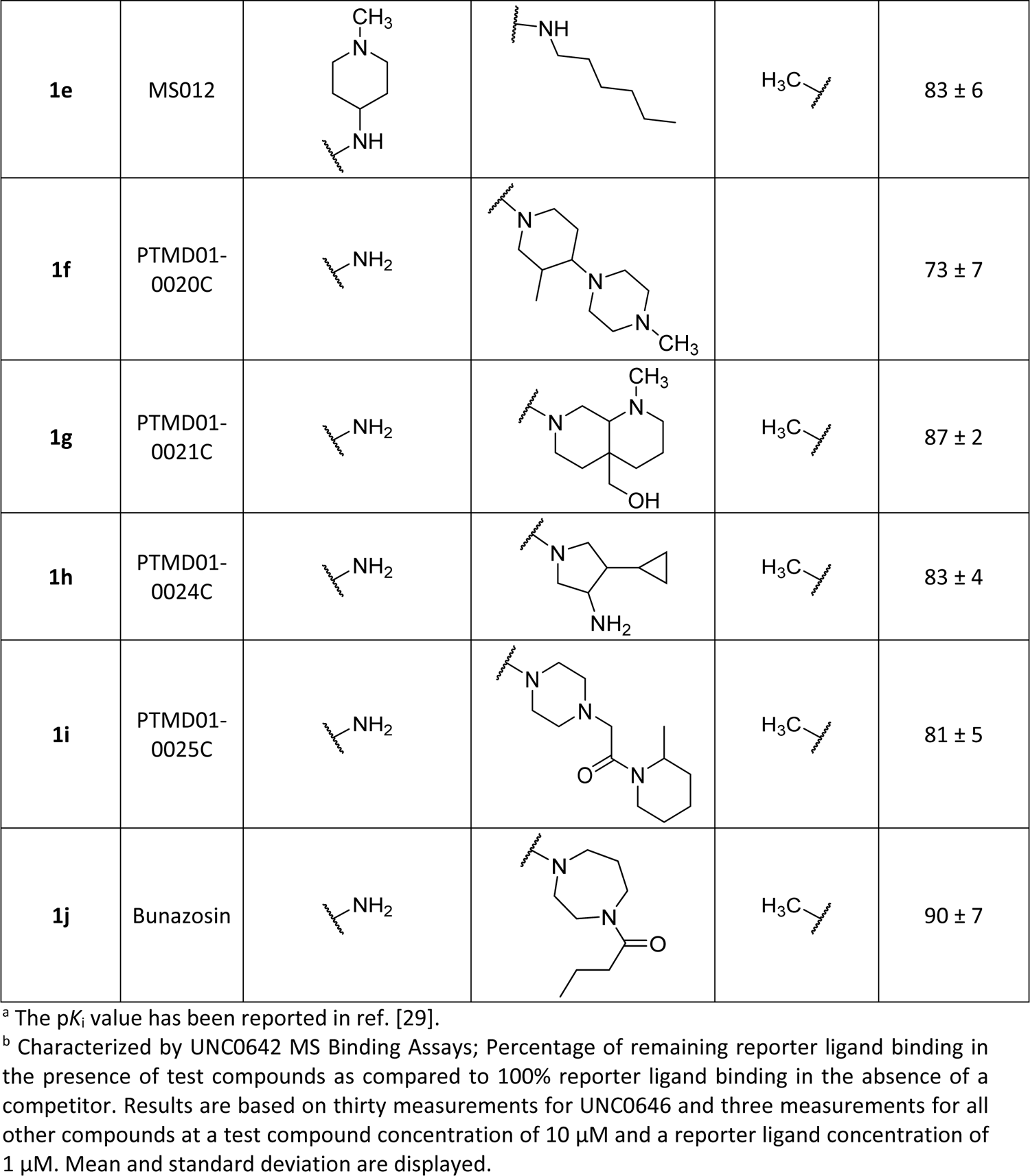
Selected analogs of UNC0646 identified by a two-dimensional similarity search and their affinities to MB327-PAM-1 in nAChR determined in MS Binding Assays.

| Compound | Name / PTMD code | R<sup>1</sup> | R<sup>2</sup> | R<sup>3</sup> | Remaining reporter ligand binding [%]<sup>b</sup> |
| --- | --- | --- | --- | --- | --- |
| **1a** | PTMD01-0019C |  |  |  | 47 ± 3 |
| **1b** | UNC0379 |  |  |  | 59 ± 4 |
| **1k** | PTMD01-0050 |  |  |  | 14 ± 0 |
| **1c** | ZT-12-037-01 |  |  |  | 66 ± 5 |
| **1d** | C-021 |  |  |  | 63 ± 2 |
| **1e** | MS012 |  |  |  | 83 ± 6 |
| **1f** | PTMD01-0020C |  |  |  | 73 ± 7 |
| **1g** | PTMD01-0021C |  |  |  | 87 ± 2 |
| **1h** | PTMD01-0024C |  |  |  | 83 ± 4 |
| **1i** | PTMD01-0025C |  |  |  | 81 ± 5 |
| **1j** | Bunazosin |  |  |  | 90 ± 7 |

<sup>a</sup> The $pK_i$ value has been reported in ref. [29].
<sup>b</sup> Characterized by UNC0642 MS Binding Assays; Percentage of remaining reporter ligand binding in the presence of test compounds as compared to 100% reporter ligand binding in the absence of a competitor. Results are based on thirty measurements for UNC0646 and three measurements for all other compounds at a test compound concentration of 10 μM and a reporter ligand concentration of 1 μM. Mean and standard deviation are displayed.

The second strongest reduction of reporter ligand binding was observed for UNC0379 (**1b**), a ligand with a substituent with increased flexibility at the 4-position compared to UNC0646. Along these lines, the results for ZT-12-037-01 (**1c**), C-021 (**1d**), MS012 (**1e**), PTMD01-0020C (**1f**), PTMD01-0021C (**1g**), PTMD01-0024C (**1h**), and PTMD01-0025C (**1i**) indicate that for affinity towards MB327-PAM-1, the positively charged amino side chain can be present at either position 2 or position 4 of the quinazoline building block. Furthermore, bunazosin (**1j**) has no basic side chains at the quinazoline ring but only the two electron donating groups in 2- and 4-positions, further indicating that the positive charge of the ligand might also be located within the heteroaromatic ring. This confirms the above observation that the location of the positive charge is not crucial for the binding of UNC0646 analogs. Based on these results, we synthesized PTMD01-0050 (**1k**), a chimera inspired by UNC0646, UNC0642, and UNC0379. The synthesis consisting of two steps started from **7** [Bernauer *et al.*, in preparation] [50] (Scheme 1). In analogy to a procedure described in the literature [79], **7** was reacted with *N*-methylpiperidine (2.0 equiv) at 150 °C under microwave irradiation for 1 h affording the quinazoline **8** with a piperidine ring in 2-position after column chromatography in good yield (78%). For the subsequent substitution of chloride in 4-position, **8** was stirred with 5-(pyrrolidin-1-yl)pentan-1-amine (2.0 equiv) in the presence of DIEA (3.0 equiv) under microwave irradiation at 160 °C for 15 min. The desired product PTMD01-0050 (**1k**) could be isolated in good yield (72%). Notably, this compound shows a higher reporter ligand displacement than UNC0646 (Table 1).

**Scheme 1:**
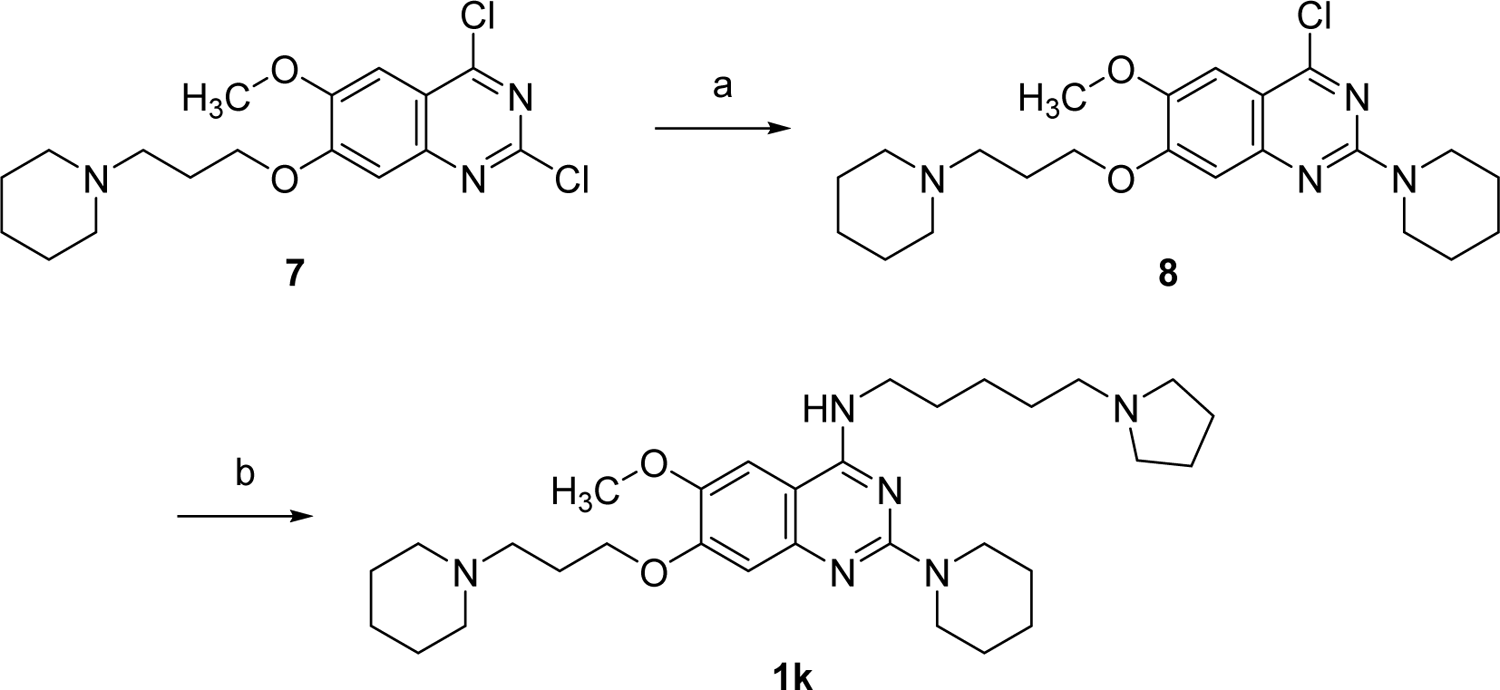
Reagents and conditions: (a) 1-methylpiperidine (2.0 equiv), 1,4-dioxane, 150 °C (300 W), 1 h, 78%; (b) 5-(pyrrolidin-1-yl)pentan-1-amine (2.0 equiv), DIEA (3.0 equiv), *i*-PrOH, microwave: 200 W, 160 °C, 15 min, 72%.

### Ligand-based screening using PTMD01-0004 (2a) as a template representing an analog of UNC0646 with a reduced molecular structure

While the two-dimensional similarity search based on UNC0646 yielded new, affine molecules binding to MB327-PAM-1, this approach did not consider the position and orientation of the ligand in MB327-PAM-1. Thus, we also performed a ligand-based screening in MB327-PAM-1 using PTMD01-0004 (**2a**) [Bernauer *et al.*, in preparation] as a template (Figure 1, yellow scheme). We started with this analog of UNC0646 because the substituent in the 2-position of UNC0646 shows minor interactions with the receptor in our proposed binding mode [29], and UNC0646 violates the molecular weight rule of Lipinski’s “rule of five” [80], in contrast to PTMD01-0004 (**2a**). Furthermore, the absence of an electron-donating group in the 2-position only has a minor impact on the affinity (Sichler *et al.*, submitted to Tox. Lett. on the 1^st^ of August 2023).

We performed a two-step screening (see Materials and Methods) using a database of synthesizable compounds based on the building blocks of PTMD01-0004 (**2a**) (SI Figure S1). We selected five compounds that we synthesized (see below) and tested for affinity towards MB327-PAM-1 (**2b**-**2e**, **2g**, Table 2). The substituents chosen for position 4 did not increase the affinity in any compound compared to PTMD01-0004 (**2a**). Still, slight modifications in this substituent can influence reporter ligand displacement significantly.

**Table 2:** Selected analogs of PTMD01-0004 (**2a**) identified by a ligand-based screening followed by template-based docking and their affinities to MB327-PAM-1 in nAChR determined in MS Binding Assays.

| Compound | PTMD code | R<sup>1</sup> | R<sup>2</sup> | Remaining reporter ligand binding [%]<sup>a</sup> |
| --- | --- | --- | --- | --- |
| **2a** | PTMD01-0004<sup>b</sup> |  |  | 50 ± 5<sup>c</sup> |
| **2b** | PTMD01-0032 |  |  | 50 ± 7 |
| **2c** | PTMD01-0053 |  |  | 76 ± 7 |
| **2d** | PTMD01-0027 |  |  | 71 ± 3 |
| **2e** | PTMD01-0030 |  |  | 60 ± 3 |
| **2f** | PTMD01-0005<sup>b</sup> |  |  | 63 ± 4<sup>d</sup> |
| **2g** | PTMD01-0043 |  |  | 43 ± 1 |

<sup>a</sup> Characterized by UNC0642 MS Binding Assays; Percentage of remaining reporter ligand binding in the presence of test compounds as compared to 100% reporter ligand binding in the absence of a competitor. If not stated otherwise, results are based on three measurements at a test compound concentration of 10 μM and a reporter ligand concentration of 1 μM. Mean and standard deviation are displayed.
<sup>b</sup> PTMD01-0004 (**2a**) [Bernauer *et al.*, in preparation] and PTMD01-0005 (**2f**) were not identified in this study but are shown as reference structures to compare to PTMD01-0043 (**2g**).
<sup>c,d</sup> Results are based on twelve and six measurements, respectively, at a test compound concentration of 10 μM and a reporter ligand concentration of 1 μM. Mean and standard deviation are displayed.

As to the UNC0646 building block, our two-dimensional similarity search revealed that substituting it with flexible linkers in the 4-position can lead to highly affine compounds as seen for PTMD01-0050 (**1k**). The compound with increased flexibility between the quinazoline ring and the basic side chain nitrogen located within the cyclohexyl ring, PTMD01-0032 (**2b**), has a higher affinity than PTMD01-0053 (**2c**). However, the piperazine ring of PTMD01-0053 (**2c**) might also result in an alternative distance between the positive charge of the side chain and the quinazoline moiety, depending on the protonation site. Still, the relation between linker flexibility and affinity is also observed in PTMD01-0027 (**2d**), PTMD01-0030 (**2e**), and PTMD01-0032 (**2b**), where the distance between the quinazoline ring to the positively charged nitrogen is 3-4 heavy atoms long. However, to verify this trend, further compound testing will be required. Additionally, this trend does not always apply. PTMD01-0053 (**2c**) with a more flexible side chain than PTMD01-0030 (**2e**) has a lower affinity. Thus, the additional polar atom and the additional methyl substituent of the piperazine ring as well as the different distance between the positive charge in the side chain and the quinazoline moiety of PTMD01-0053 (**2c**) may also lead to a decrease in affinity.

In the 7-position, we identified in PTMD01-0043 (**2g**), an alternative substituent that leads to a higher reporter ligand displacement than if the same substituent as in UNC0646 is used in the in 7-position [PTMD01-0005 (**2f**)]. PTMD01-0043 (**2g**) otherwise bears the same side chains as PTMD01-0005 (**2f**) except in the 7-position of the quinazoline ring. As for the assessment of the two-dimensional similarity search, we verified our results by characterizing the binding affinity of the most affine compound according to the single point determinations [PTMD01-0043 (**2g**); remaining reporter ligand binding 43 ± 1%] in a full-scale MS Binding Assay yielding a p*K*_i_ of 5.46 ± 0.04 (SI Figure S6).

#### Synthesis of compounds 2b-g

Target compounds **2b-2f** were easily accessible by a two-step synthesis from commercially available building block **11** (Scheme 2). First, key intermediate **9** [51] was obtained in quantitative yield (> 99%) by reaction of quinazoline derivative **11** with 1.25 equiv 3-(piperidin-1-yl)propan-1-ol under *Mitsunobu* conditions (1.3 equiv PPh_3_, 1.3 equiv DBAD, THF, rt, 20 h) following a literature procedure [81]. In the second step, the 4-amino substituents were introduced to afford the target compounds **2b-2f**. Nucleophilic displacement of the 4-chloro substituent was achieved according to a procedure reported in the literature [82] by heating **9** with the corresponding amines (2.0 equiv) in the presence of DIEA (3.0 equiv) to 160 °C under microwave irradiation. Thus, 4-aminoquinazolines **2b-2d** and **2f** were isolated in good to excellent yields (78-93%). However, the reaction with the sterically demanding amine 1-(azepan-1-yl)-2-methylpropan-2-amine to get **2e** was sluggish. Hence, a higher excess of the amine (10 equiv) was applied. This led to the target compound, which could be isolated in a yield of 25% only, which is partly due to the fact, that also a small amount of a side-product had formed being difficult to separate.

**Scheme 2:**
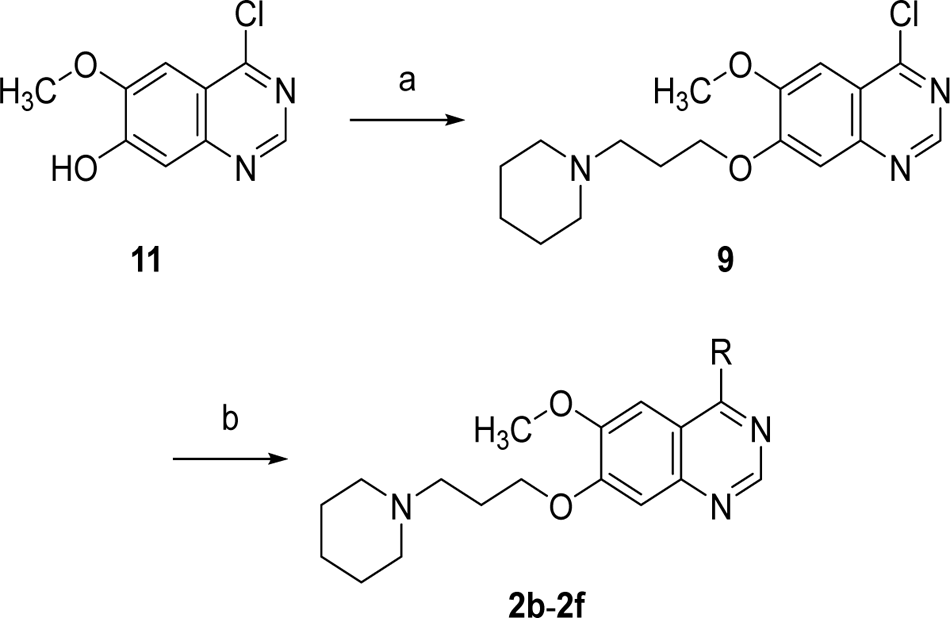
Reagents and conditions: (a) 3-piperidin-1-ylpropan-1-ol (1.25 equiv), PPh_3_ (1.3 equiv), DBAD (1.3 equiv), THF, rt, 20 h, > 99%; (b) amines (2.0-10 equiv), DIEA (3.0 equiv), *i*-PrOH, microwave: 200 W, 160 °C, 15 min-60 min, **2b**: 93%, **2c**: 78%, **2d**: 89%, **2e**: 25%, **2f**: 93%.

The 7-aminoquinazoline **2g** was synthesized in two steps starting from commercially available 7-fluoro substituted quinazoline-4(3*H*)-one **12** (Scheme 3). In the first step, the lactame **12** was converted to the 4-aminoquinazoline **10** by a phosphonium-mediated S_N_Ar reaction according to a procedure described in the literature [83]. Thus, **12** was reacted with 1.5 equiv of 1-propan-2-ylpiperidin-4-amine, PyBOP and DBU in acetonitrile at rt for 1 h, to obtain product **10** in excellent yield (92%). The subsequent substitution of the fluorine in 7-position of **10** to afford target compound **2g** was achieved by a reaction of **10** with a 5-fold excess of 1-(pyrrolidin-3-ylmethyl)piperidine in NMP at 135 °C in the presence of K_2_CO_3_ (1.1 equiv) according to a procedure reported in the literature [84]. In this way, the product **2g** could be isolated in 90% yield and high purity (99%).

**Scheme 3:**
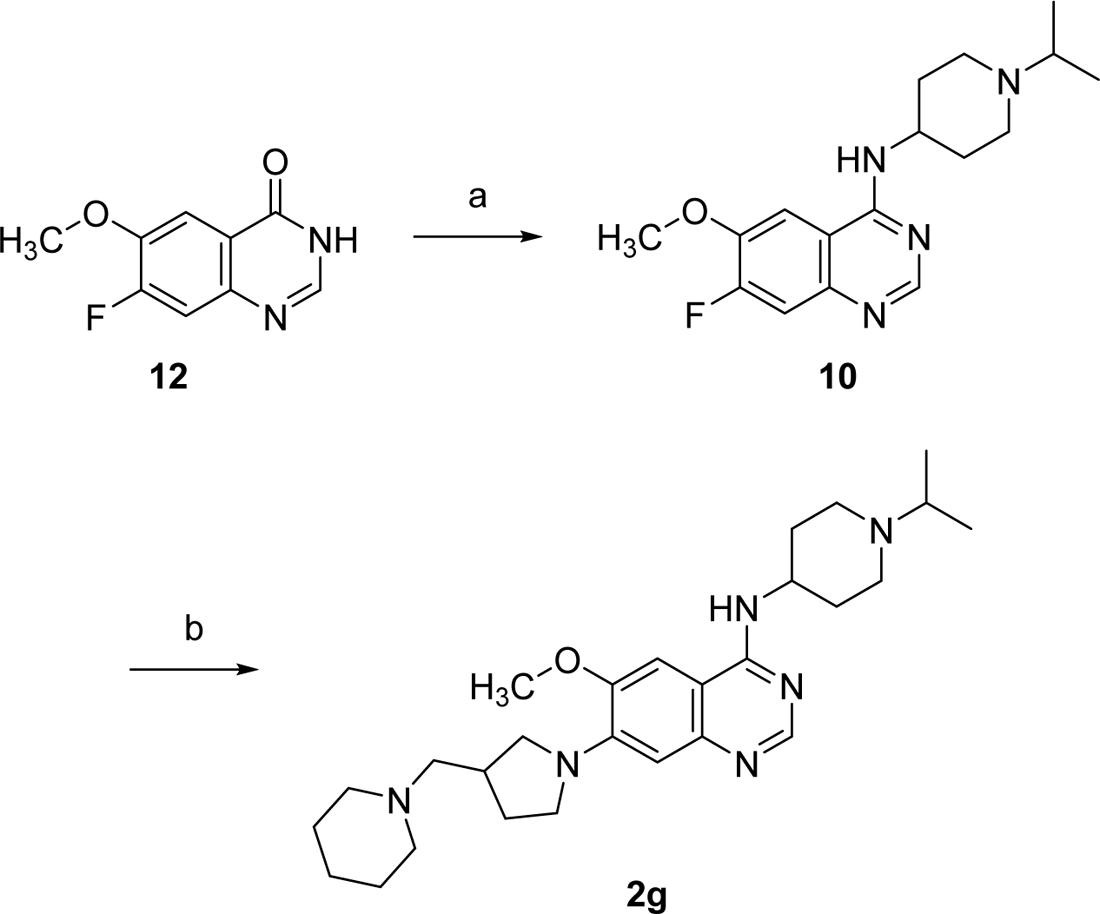
Reagents and conditions: (a) 1-propan-2ylpiperidin-4-amine (1.5 equiv), PyBOP (1.5 equiv), DBU (1.5 equiv), acetonitrile, rt, 1 h, 92%; (b) 1-(pyrrolidin-3-ylmethyl)piperidine (5.0 equiv), K_2_CO_3_ (1.1 equiv), NMP, 135 °C, 20 h, 90%.

### Structure-based screening reveals new chemotypes with a higher affinity than MB327

We first screened the lead-like library of ZINC20 [53] with 3,434,621 molecules using the homology model of the human muscle-type nAChR and OpenEye FRED [54-56] as docking engine with default parameters (Figure 1, green scheme). However, we know from previous work that larger molecules, such as UNC0646, usually bind to MB327-PAM-1 with a higher affinity than smaller ones, such as MB327. Furthermore, the two previously identified binders in MB327-PAM-1, UNC0646, and MB327, carry at least two positive charges. Thus, we decided to also screen a subset of the ZINC20 database [53] containing all doubly protonated in-stock compounds (129,606 compounds) by docking into the *Torpedo* nAChR (see also above).

We ordered 12 compounds based on visual inspection of the best 1000 hits in MB327-PAM-1 in each subunit in each of the screenings (2 x 5 x 1000 = 10,000 hits in total) (Figure 2, SI Table S3) (PTMD99-0001C – PTMD99-0015C). (In preliminary MS binding studies (the results of which had later on to be partly revised; for final results see SI Table S3), PTMD99-0006C (**13**), PTMD99-0010C (**14**), and PTMD99-0014C (**15**) showed the most promising results.) Thus, we decided to inspect the best 1000 hits in both screenings in each subunit again to find structurally similar chemotypes. We ordered three analogs of PTMD099-0006C (**13**), seven analogs of PTMD99-0010C (**14**), and eight analogs of PTMD99-0014C (**15**) (SI Table S4). In each group, at least one compound (at 10 μM concentration) displaced the reporter ligand UNC0642 (at 1 μM concentrations) from MB327-PAM-1 during single-concentration MS Binding Assay experiments indicating that these compounds show a higher affinity towards MB327-PAM-1 than MB327, which shows a remaining marker ligand binding of 102 ± 9% (*n* = 6) under identical conditions (1 μM reporter ligand, 10 μM test ligand) (Bernauer *et al.*, in preparation). In total, four new chemotypes, all containing at least one positive charge, were identified that displace UNC0642 from MB327-PAM-1 at concentrations of 10 μM (reporter ligand concentration of 1 μM) to any appreciable extent. However, for two of these compounds the remaining reporter ligand binding values are scarcely not significantly different from 100%, while the two other compounds differ significantly from 100% {*p* < 0.05 according to a two-sided one-sample *t*-test; *p* [PTMD99-0001C (**3**)] = 0.064, *p* [PTMD99-0016C (**4**)] = 0.005, *p* [PTMD99-0026C (**5**)] = 0.079, *p* [cycloguanil (**6**)] = 0.006}.

**Figure 2:**
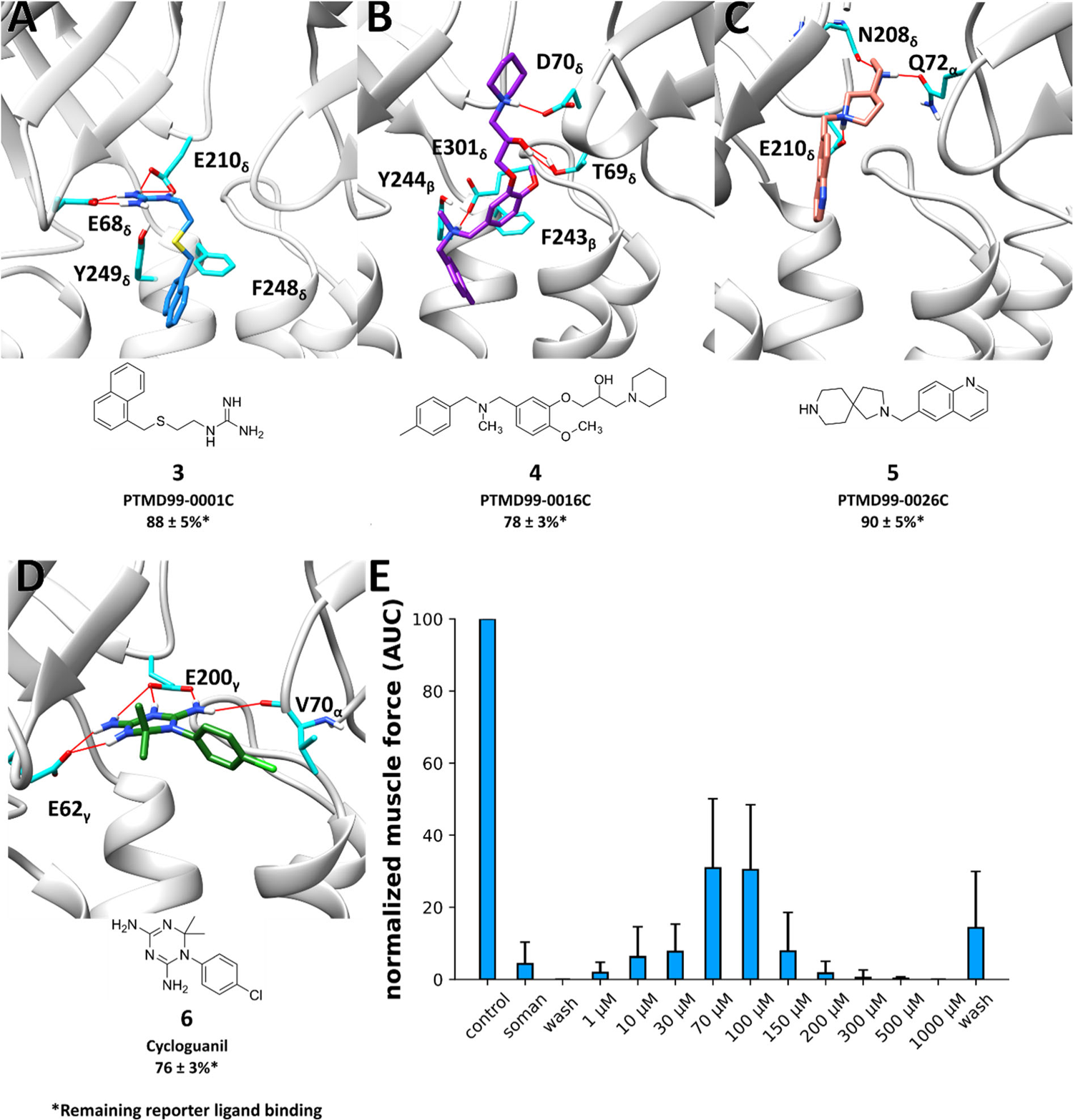
Docked binding mode and MS Binding Assay affinity data of selected hits from a structure-based screening in different subunits and species of nAChR. Docked binding mode of **(A)** PTMD99-0001C (**3**) in between the α- and δ-subunits of the human nAChR, **(B)** PTMD99-0016C (**4**) in between the δ- and β-subunits of the *Torpedo* nAChR, **(C)** PTMD99-0026C (**5**) in between the α- and δ-subunits of the *Torpedo* nAChR, and **(D)** cycloguanil (**6**) in between the α- and γ-subunits of *Torpedo* nAChR. Red lines indicate hydrogen bonds. Percentage values indicate the remaining reporter ligand binding in the presence of test compounds (at 10 μM concentration) as compared to 100% reporter ligand binding in the absence of a competitor using the reporter ligand UNC0642 in MS Binding Assays (1 µM UNC0642) (mean ± SD, *n* = 3). Compounds displaying chirality were tested as racemats. **(E)** Resoration of muscle force of soman-inhibited muscles after treatment with cycloguanil (**6**). Error bars indicate the standard deviation (*n* is between 5 and 27). Since the largest efficacies are observed at low stimulation frequencies [10], results are only shown for a stimulation frequency of 20 Hz (see SI Table S5 for all stimulation frequencies applied).

Of the analogs based on PTMD99-0006C (**13**), PTMD99-0016C (**4**) shows the highest affinity within this group and is the only compound able to displace UNC0642 to any appreciable extent during measurements with test compound concentrations of 10 μM and reporter ligand concentrations of 1 µM. Small changes in the 4-methylbenzyl group can have a high impact on affinity. For example, PTMD99-0020C (**16**) (SI Table S4), bearing a (3-methylpyridin-4-yl)methyl substituent instead of the 4-methylbenzyl group, does not show a displacement of the reporter ligand to any appreciable extent anymore under identical experimental conditions. In fact, all compounds bearing a heteroaromatic ring instead of the 4-methylbenzyl group fail to displace UNC0642 to any appreciable extent under identical experimental conditions to a reasonable extent.

Based on the initial results for PTMD99-0010C (**14**), we identified PTMD99-0026C (**5**), able to displace the reporter ligand (concentration 1 μM) to any appreciable extent at 10 μM test compound concentration. However, compounds with an amide group in 3-position to the nitrogen at position 2 of the 2,8-diazaspiro[4.5]decane system do not displace the reporter ligand UNC0642 under similar experimental conditions to any appreciable extent (PTMD99-0010C (**14**), −0023C (**17**), −0024C (**18**), - 0025C (**19**), −0028C (**20**) (SI Table S3, S4)). Furthermore, replacing the quinolinyl substituent by a 5-(*tert*-butyl)-pyrazol-3-yl substituent (PTMD99-0031C (**21**) (SI Table S4)) also abrogates the reporter ligand displacement indicating that hydrogen bond donors as substituents of the 2,8-diazaspiro[4.5]decane ring might be unfavorable.

Additionally, as the fourth novel chemotype binding to MB327-PAM-1, the 1,6-dihydro-1,3,5-triazine-2,4-diamine building block was identified. Most interesting, the compound showing the highest affinity, cycloguanil (**6**, 10 μM test compound concentration at 1 μM reporter ligand concentration, Figure 2D), is the active metabolite of the antimalarial drug proguanil. In competitive MS binding experiments, we observed a p*K*_i_ value for cycloguanil (**6**) of 3.64 ± 0.03 (SI Figure S7), significantly higher compared to MB327 [p*K*_i_ (MB327) = 3.40 ± 0.04 [29], *p* < 0.01, according to a two-sided *t*-test]. Cycloguanil (**6**) forms salt bridges both with E62_γ_ and E200_γ_ in the docked pose (Figure 2D). These two amino acids are highly conserved among different subunits of several species (Table 3), including the *Torpedo* nAChR, which is used in our MS Binding Assay, the rat muscle nAChR, which is used in our rat diaphragm assays, and in the human nAChR, in which the compounds need to exhibit an effect after OPC poisoning. Furthermore, these glutamates are crucial for the stabilization of the calcium ion in the α7 nAChR that can act as a positive allosteric modulator [85-87]. According to our screening results, we can, in general, see that larger substituents at both rings of cycloguanil (**6**) lead to a decrease in affinity (**22-28**, SI Table S4). Compounds based on the 1,3,5-triazin-2,4-diamin building block are overall much smaller than UNC0646 (M [UNC0646] = 621.93 g mol^-1^; M [Cycloguanil (**6**)] = 251.72 g mol^-1^), leading to compounds with an improved ligand efficiency.

**Table 3:** Sequence conservation of E62_γ_ and E200_γ_ (green shadings) in the human muscle-type, *Torpedo*, and rat nAChR with respect to structurally homologous positions in the γ-subunit of the *Torpedo* nAChR.^a^.

| Human muscle type |  |  |  | *Torpedo* |  |  |  | Rat |  |  |  |
| --- | --- | --- | --- | --- | --- | --- | --- | --- | --- | --- | --- |
| α | β | δ | ε | α | β | δ | γ | α | β | δ | ε |
| E | E | E | E | E | E | E | E62 | E | E | E | E |
| E | Q | E | E | E | Q | E | E200 | E | Q | E | E |

<sup>a</sup> E62<sub>γ</sub> and E200<sub>γ</sub> are important for interactions with cycloguanil (**6**) in the docked binding mode and during MD simulations (Figure 3; see also text).

Enough substance to conduct competition experiments with varying ligand concentrations and to perform rat diaphragm assays in order to investigate the restoration of muscle force after soman poisoning was only commercially available for cycloguanil (**6**). Treatment with cycloguanil (**6**) led to significant restoration of muscle force in rat diaphragm hemispheres after soman inhibition (Figure 2E, SI Table S5). The maximum restoration at stimulation frequencies of 20 Hz is comparable to the maximum restoration when using MB327 as a treatment option. However, while concentrations of 300 µM are necessary for the maximum effect of MB327 [26.29 ± 18.43% (mean ± SD; *n* = 27) restoration of muscle force, values taken from ref. [13]], cycloguanil (**6**) exerts a comparable effect at concentrations of 70 μM (30.87 ± 19.23%; *n* = 5). At a concentration of 100 µM, cycloguanil (**6**) leads to a significantly increased restoration of muscle force compared to MB327 [30.42 ± 18.04% vs. 17.77 ± 7.5%, values for MB327 taken from ref. [13], *p* < 0.01 according to a two-sided *t*-test (*n* = 27)]. Like MB327, cycloguanil (**6**) has a small therapeutic index, leading to muscle force inhibition at concentrations ≥ 300 μM (Figure 2E, SI Figure S8). Thus, cycloguanil (**6**) currently cannot be considered as a treatment option but as a novel lead structure for treating OPC poisoning.

To further investigate the binding mode of cycloguanil (**6**), we performed MD simulations starting from the docked conformation. In 6 out of 10 replicas over 1 μs simulation time each, the ligand left the binding site (SI Figure S9). In the replicas where cycloguanil (**6**) remained in the binding site, the binding mode shifted. Whereas the interaction with the two glutamates persisted, the aromatic system of cycloguanil (**6**) moved towards the transmembrane region of nAChR in the direction of Y239_γ_ (Figure 3A). This amino acid is located in a hydrophobic part of the binding site. Thus, we clustered the replica in which cycloguanil (**6**) remained in the binding site and performed additional 10 replicas of 1 μs long MD simulations starting from a representative structure. During the simulations, the membrane and receptor remained structurally virtually invariant (SI Figure S10, S11). Cycloguanil (**6**) continued to remain in the binding site in all replicas and showed highly conserved interactions with E62_γ_ and E200_γ_ (Figure 3B, C). Thus, we conclude that according to the MD-optimized binding mode, the interactions with the two glutamates persist and the hydrophobic interactions with amino acids close to Y239_γ_ are important for ligand stabilization.

**Figure 3:**
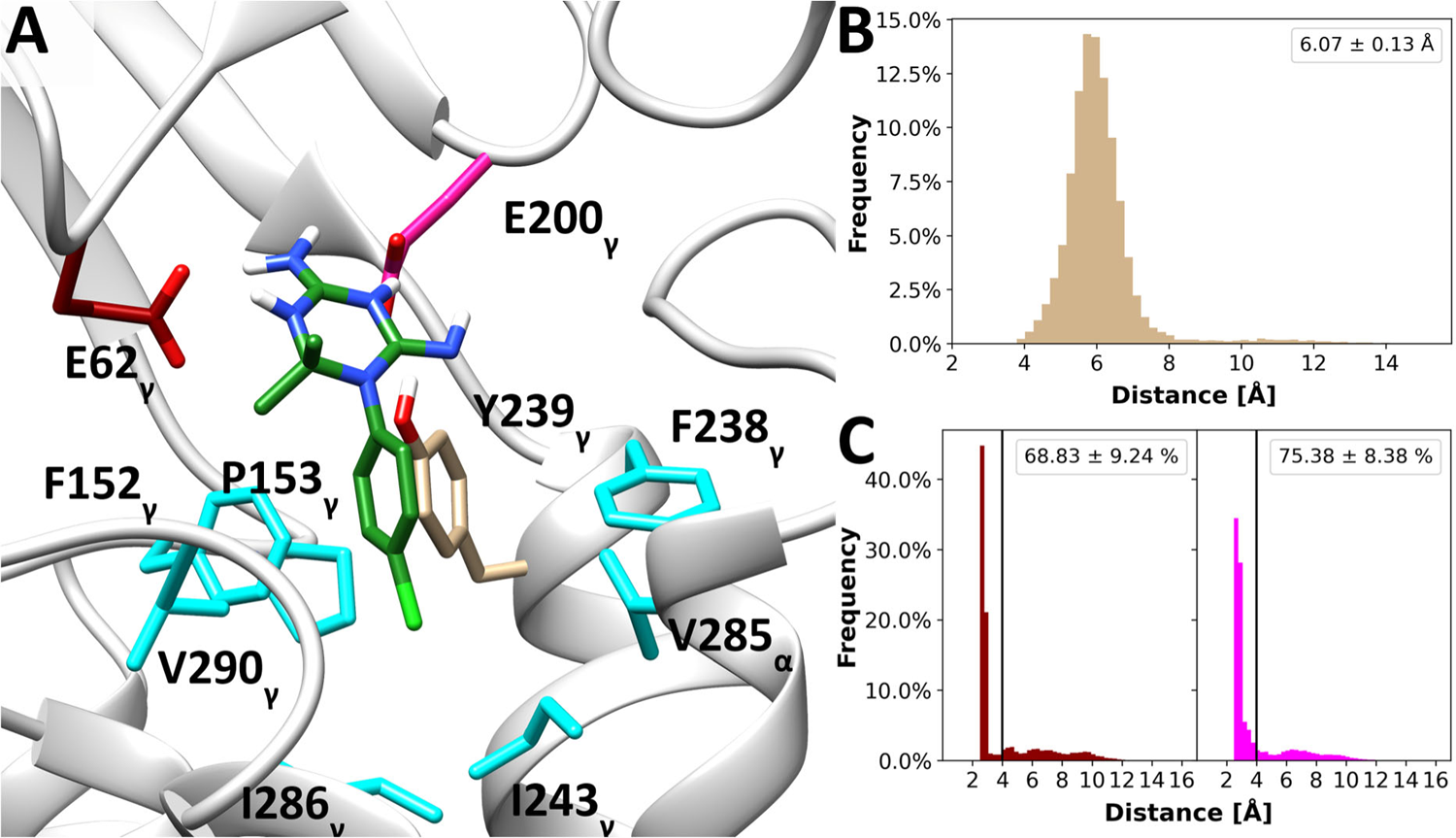
Binding mode of cycloguanil during MD simulations. **A)** Representative (according to a *k-means* clustering based on receptor and ligand atoms; the biggest cluster containing 48.2% of all frames is shown) binding mode of cycloguanil during 10 replicas of 1 μs long unbiased MD simulations starting from the docked conformation. **B)** Distance of the center of mass (COM) of the phenyl ring of cycloguanil to the phenyl ring of Y239_γ_. The mean ± SEM distance is displayed as a legend. **C)** Distance of the nitrogens that can act as hydrogen bond donors of cycloguanil to the side chain oxygens of E62_γ_ (dark red) and E200_γ_ (pink). The frequency of contacts (distance < 4 Å; mean ± SEM) is displayed as a legend.

### Prediction of pharmacokinetic and toxicological properties of best hits

UNC0646, the best hits of both ligand-based screenings [PTMD01-0050 (**1k**), PTMD01-0043 (**2g**)], and the best hit of each novel chemotype from the structure-based screening [PTMD99-0001C (**3**), PTMD99-0016C (**4**), PTMD99-0026C (**5**), and cycloguanil (**6**)] were initially probed in a pan interference compounds (PAINS) filter as implemented in the PAINS-remover webserver [71]; all compounds passed this filter, suggesting that they are less likely to react nonspecifically with biological targets. We further predicted the pharmacokinetic and toxicological properties using Schrödinger QikProp [70] and NEXUS Derek [72] (Table 4, Table 5). By far the most predictions for UNC0646 fall outside a 95% range for values of known drugs, questioning the drug-like properties of this compound. Along these lines, UNC0646 shows the worst Caco-2 cell permeability prediction as a model for gut-blood barrier permeation among all tested compounds and also violates two rules of Lipinski’s rule of five and one rule of Jorgensen’s rule of three, which are used as indicators for oral bioavailability. By contrast, all newly identified chemotypes show no violations of Lipinski’s rule of five, and only PTMD99-0026C (**5**) violates one rule of Jorgensen’s rule of three. Note, however, that the violated pharmacokinetic descriptors describe oral availability, whereas in the case of OPC poisoning drugs may be injected. On the other hand, improved oral bioavailability and reduced side effects might lead to the possibility to provide the antidote to a broader group of civilians and military members in the case of a high risk of OPC poisoning. Finally, the newly identified chemotypes [PTMD99-0001C (**3**), PTMD99-0016C (**4**), PTMD99-0026C (**5**), and cycloguanil (**6**)] show a reduced predicted affinity towards the HERG K^+^ channel and a reduced toxicological alert count compared to UNC0646 and its analogs [PTMD01-0050 (**1k**), PTMD01-0043 (**2g**)]. Also, for the new chemotypes – except PTMD99-0026C (**5**) – no bacterial mutagenicity is predicted. In that respect, all novel compounds identified from the screenings show improved predicted pharmacokinetic properties compared to UNC0646 (Table 4) and, besides PTMD99-0026C (**5**), all novel chemotypes also display improved predicted toxicological properties (Table 5).

**Table 4:**
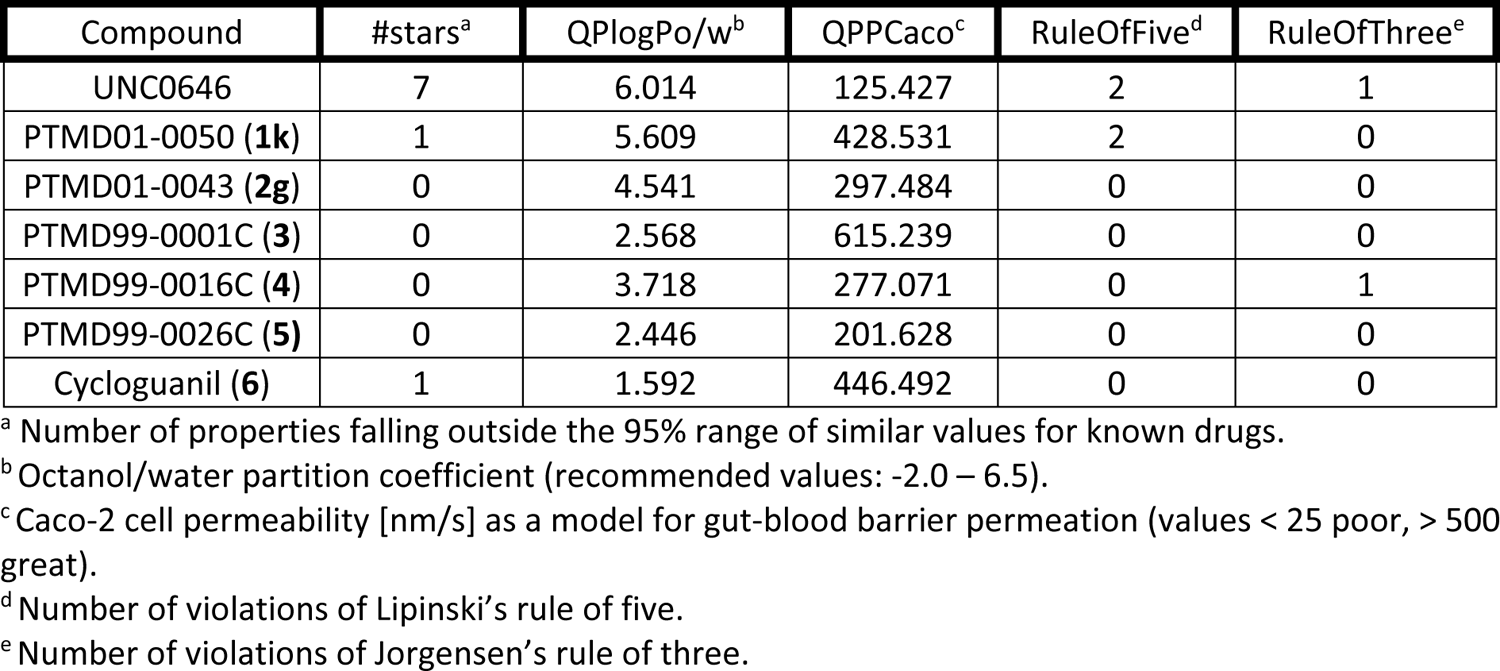
Predicted pharmacokinetic properties of the best screening hits.

**Table 5:** Predicted toxicological properties of the best screening hits.

| Compound | QPlogHERG<sup>a</sup> | Toxicological alert count<sup>b</sup> | Bacterial mutagenicity<sup>d</sup> |
| --- | --- | --- | --- |
| UNC0646 | -8.224 | 4 | EQUIVOCAL |
| PTMD01-0050 (**1k**) | -6.109 | 2 | EQUIVOCAL |
| PTMD01-0043 (**2g**) | -7.460 | 4 | EQUIVOCAL |
| PTMD99-0001C (**3**) | -4.980 | 0 | INACTIVE |
| PTMD99-0016C (**4**) | -6.125 | 2 | INACTIVE |
| PTMD99-0026C (**5**) | -6.528 | 4 | PLAUSIBLE |
| Cycloguanil (**6**) | -3.989 | 1 | INACTIVE |

<sup>a</sup> IC<sub>50</sub> value for blockage of HERG K<sup>+</sup> channels (recommended values: above -5).
<sup>b</sup> Number of alerts for toxicological predictions.
<sup>c</sup> Bacterial mutagenicity *in vitro*.

## Conclusion

To find new compounds representing novel chemotypes that bind to MB327-PAM-1 and to better understand structure-affinity relationships of the known binder UNC0646, we performed exhaustive virtual screening followed by an MS Binding Assay. As to the importance of the substituents of UNC0646 analogs, overall, beneficial substituents in position 4 are also more flexible, suggesting that conformational adaptability may be favorable compared to the loss of conformational entropy. Furthermore, while all compounds known to bind to MB327-PAM-1 carry at least one positive charge, our results indicate that the location of the positive charge plays a minor role. Based on our results, we developed PTMD01-0050 (**1k**), which leads to a higher reporter ligand displacement at test compound concentrations of 10 μM than UNC0646.

UNC0646 analogs in general show increased binding affinity with increased molecular weight and size. Due to concerns for oral bioavailability and because for some pharmacokinetic and toxicological predictions UNC0646 lies outside the recommended value range, we also aimed to find novel chemotypes binding to MB327-PAM-1. Identified compounds with four novel chemotypes can displace UNC0642 from MB327-PAM-1 (mean ± SD < 100%) at test compound concentrations of 10 μM and reporter ligand concentrations of 1 μM. While one compound (PTMD99-0016C (**4**)) already has a molecular weight > 400 Da, the other three hits have a molecular weight < 300 Da. These compounds can be used as a starting point for optimization in terms of affinity, pharmacokinetics, and resensitization capability of a desensitized nAChR. One of those compounds, cycloguanil (**6**), was tested for its resensitizing capabilities in soman-inhibited rat muscles and leads to a significantly increased restoration of muscle force compared to MB327 at a concentration of 100 μM.

The identification of more potent resensitizers of nAChR is of utmost importance to improve the currently insufficient treatment after OPC poisonings. Identifying novel chemotypes by structure-based screening and showing with our MS Binding Assay that these compounds can bind in the same binding site as MB327 suggests that the hits also bind to the allosteric binding site MB327-PAM-1.

## Acknowledgments

This work was supported by the German Ministry of Defense (E/U2AD/KA019/IF558). We are grateful for computational support and infrastructure provided by the “Zentrum für Informations-und Medientechnologie” (ZIM) at the Heinrich Heine University Düsseldorf and the computing time provided by the John von Neumann Institute for Computing (NIC) to HG on the supercomputer JUWELS at Jülich Supercomputing Centre (JSC) (user IDs: VSK33, nAChR). HG is grateful to OpenEye Scientific Software for granting a Free Public Domain Research License.

## Data availability

Data will be made available on request.

## Author contribution

J.K. performed modeling, screening, and MD simulation experiments. C.G. supported the computational experiments. T.B. synthesized analogs of UNC0646. V.N. performed MS Binding Assays, and T.S. performed rat diaphragm assays. H.G. conceived the study and supervised the project. G.H., K.N., F.P., K.W., D.S., and F.W. supervised respective study parts. All authors contributed to writing the manuscript.

