## Supporting Information for "Identification of ligands binding to MB327-PAM-1, a binding pocket relevant for resensitization of nAChRs"

### Table of Content

Supplemental Figures and Tables

Supplemental Figure S1 ………….. 3

Supplemental Figure S2 ………….. 4

Supplemental Figure S3 ………….. 5

Supplemental Figure S4 ………….. 6

Supplemental Figure S5 ………….. 7

Supplemental Figure S6 ………….. 7

Supplemental Figure S7 ………….. 7

Supplemental Figure S8 ………….. 8

Supplemental Figure S9 ………….. 9

Supplemental Figure S10 ………….. 9

Supplemental Figure S11 ………….. 10

Supplemental Table S1 ………….. 11

Supplemental Table S2 ………… 12-13

Supplemental Table S3 ………… 14-15

Supplemental Table S4 ………… 16-17

Supplemental Table S5 ………… 18

Analytical Data ………… 19-21

Supplemental References ………….. 22

### Supplemental Figures

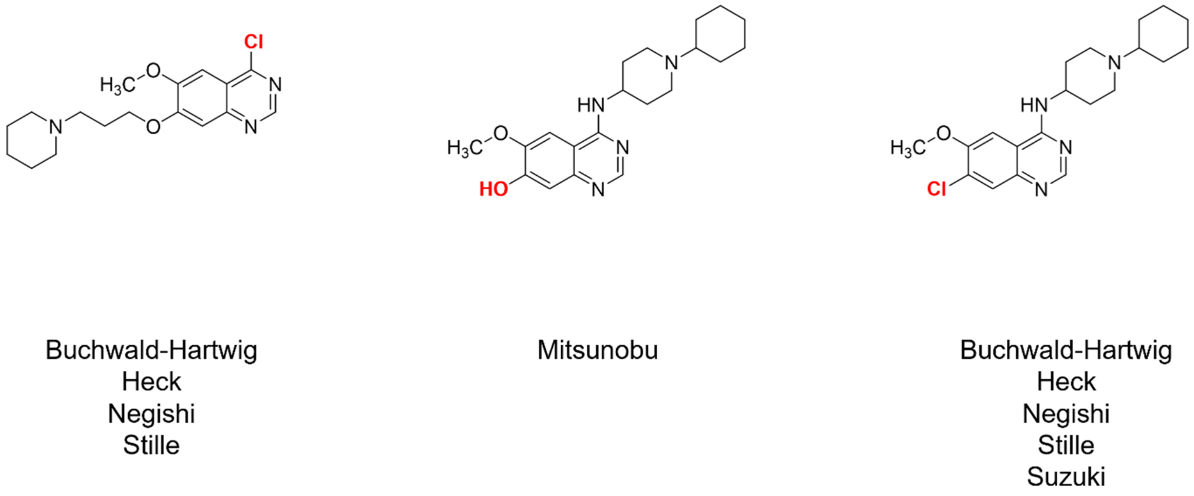


**Figure S1:** Building blocks of PTMD01-0004 (**2a**) for the generation of a virtual database. For each building block, the stated virtual syntheses have been performed with building blocks available on MolPort (https://molport.com) using PINGUI [1]. Functional groups that participate in the respective reaction are shown in red.

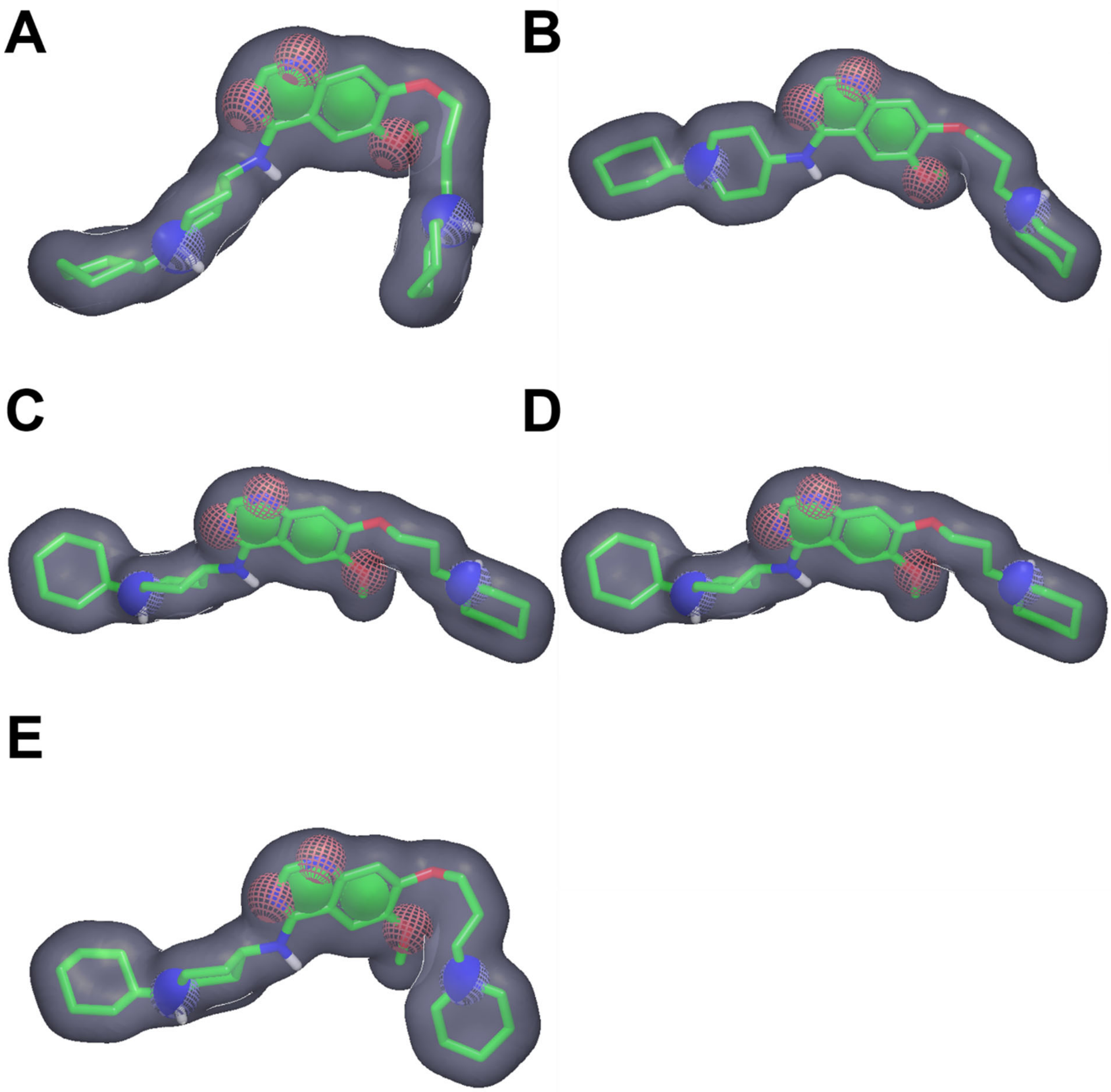


**Figure S2:** Pharmacophore models used for database filtering during template-based docking. Conformations of PTMD01-0004 (**2a**) (green) are shown as sticks between the **A)** α- and δ-, **B)** δ- and β-, **C)** β- and α-, **D)** α- and γ-, and **E)** γ- and α-subunits. Colored elements indicate pharmacophore filters: the grey surface indicates the ligand surface, green circles indicate aromatic systems, blue circles indicate hydrogen bond donor cations, and red circles indicate hydrogen bond acceptors. Figures were generated using OpenEye vROCS [2].

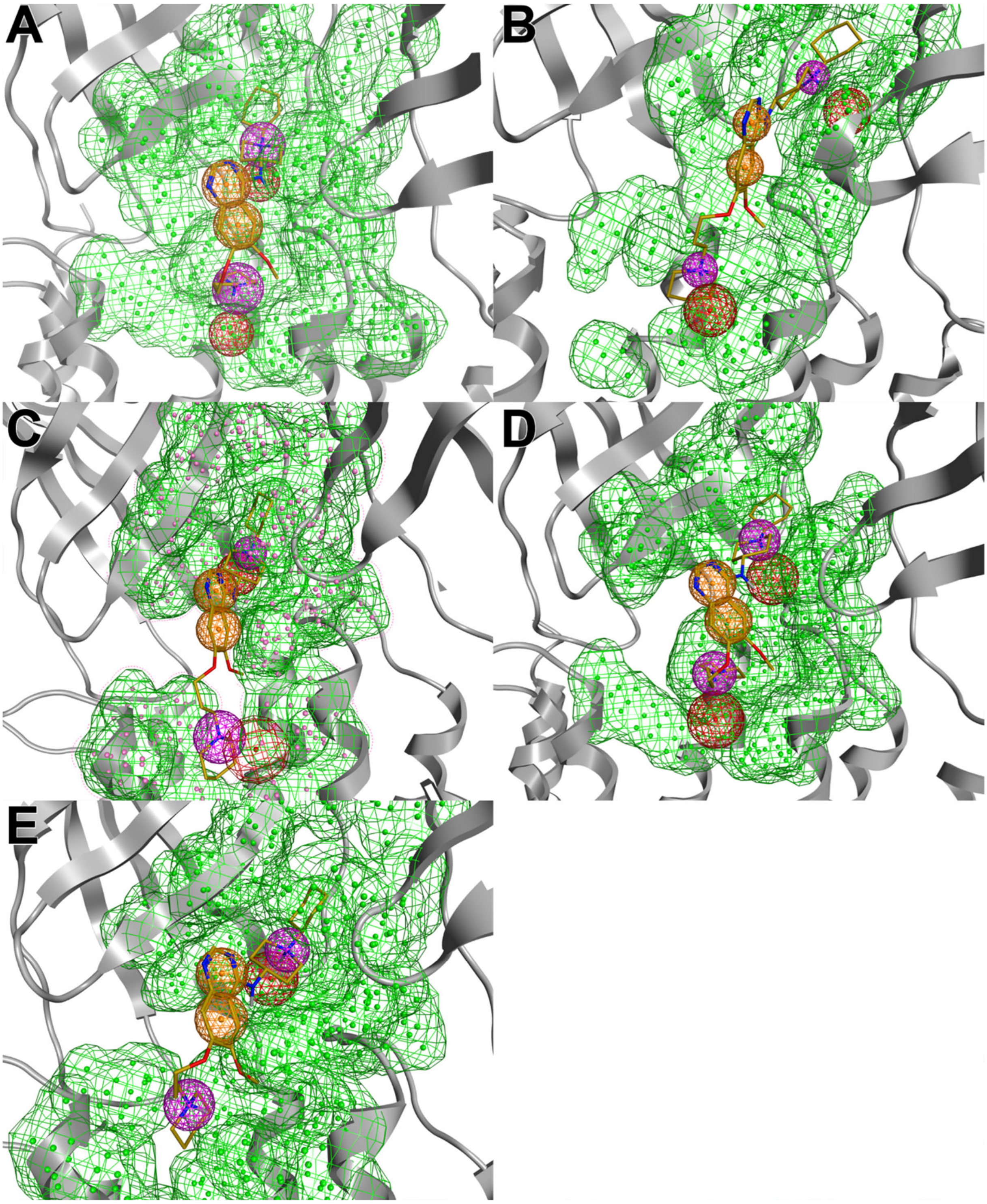


**Figure S3:** Binding mode of and pharmacophore model for template-based docking based on PTMD01-0004 (**2a**) (gold) shown between the **A)** α- and δ-, **B)** δ- and β-, **C)** β- and α-, **D)** α- and γ-, and **E)** γ- and α-subunits. Colored mesh areas indicate features of the pharmacophore filter: green area indicates the surface of the receptor where no atoms of the ligands are allowed to overlap, orange circles indicate the presence of an aromatic system, purple circles indicate the presence of a cation donor, and red circles indicate the direction of the hydrogen bond donor orientation. Figures were generated using Chemical Computing Group MOE [3].

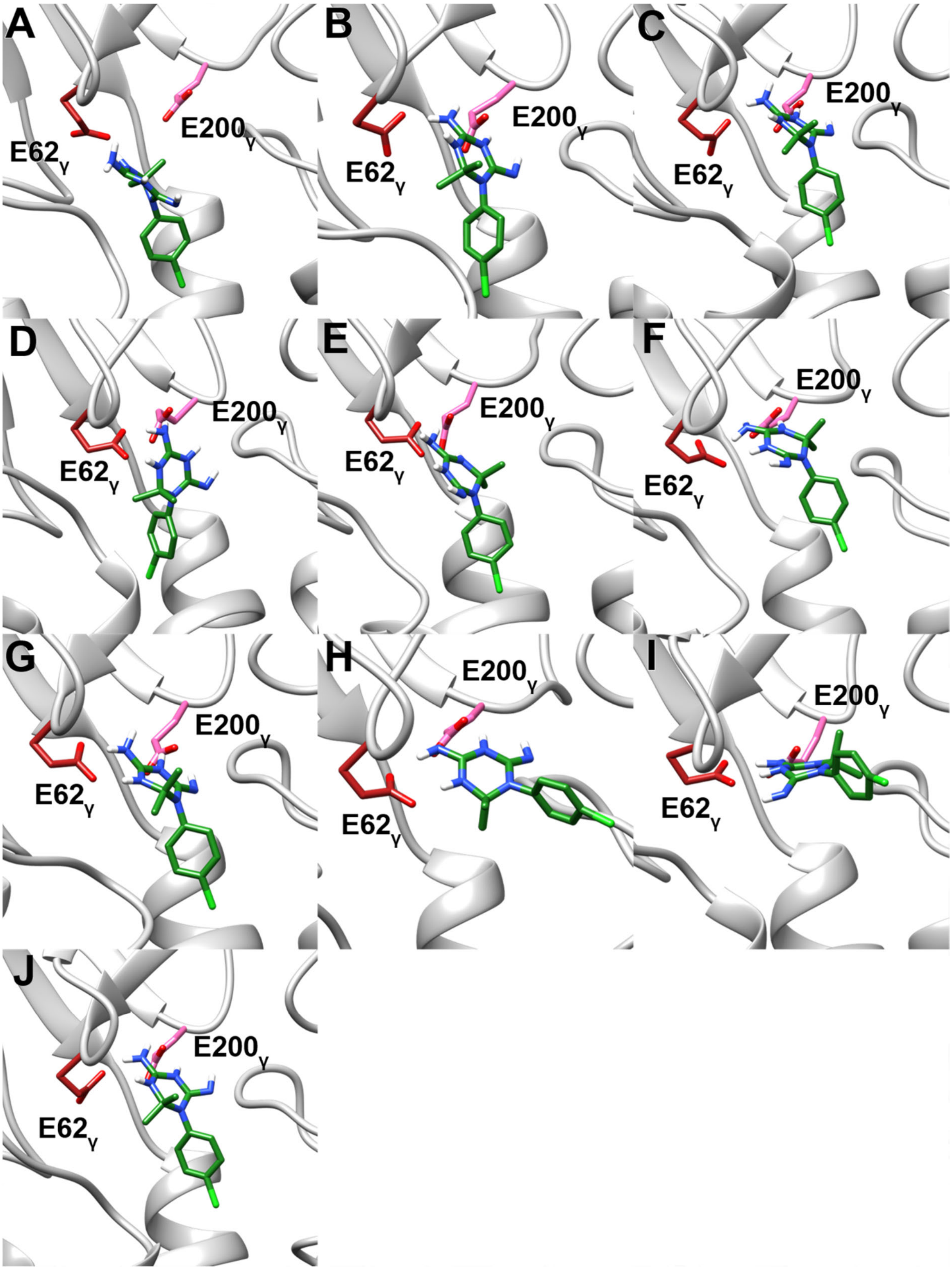


**Figure S4:** Representative binding modes of cycloguanil in MD simulations starting from the docked binding mode after clustering using the *k-means* algorithm as implemented in CPPTRAJ [4]. **A)** Only in the largest cluster (containing 18.3% of all frames) cycloguanil is not interacting with E200$_{\gamma}$. In the **B)** second (containing 14.1% of all frames), **C)** third (containing 12.4 % of all frames, **D)** fourth (containing 10.9% of all frames), **E)** fifth (containing 10.8% of all frames), **F)** sixth (containing 10.5% of all frames), **G)** seventh (containing 10.1% of all frames), **H)** eight (containing 6.7% of all frames), **I)** ninth (containing 3.6% of all frames), and **J)** tenth (containing 2.5% of all frames) largest cluster, cycloguanil is interacting with E200$_{\gamma}$ and E62$_{\gamma}$.

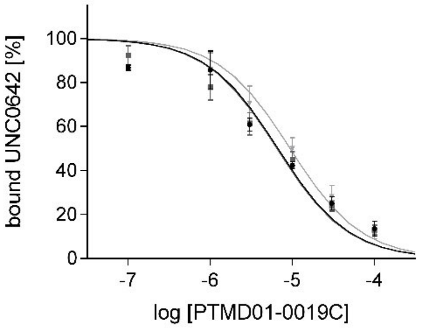


**Figure S5:** Competition curves obtained for PTMD01-0019C (**1a**) in UNC0642 MS Binding Assays. Data points (mean ± SD, *n* = 3) represent the specific binding of UNC0642.

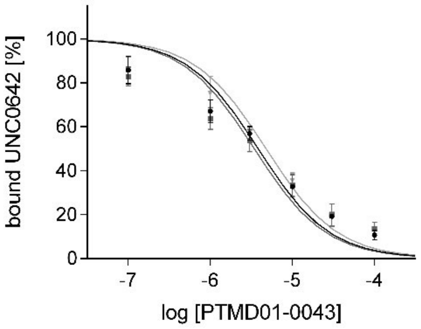


**Figure S6:** Competition curves obtained for PTMD01-0043 (**2g**) in UNC0642 MS Binding Assays. Data points (mean ± SD, *n* = 3) represent the specific binding of UNC0642.

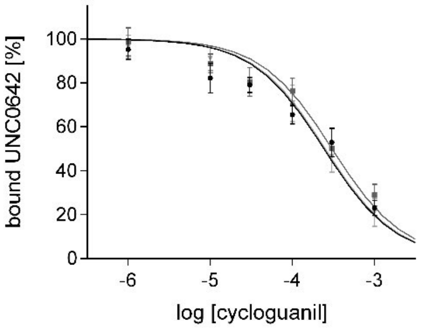


**Figure S7:** Competition curves obtained for cycloguanil (**6**) in UNC0642 MS Binding Assays. Data points (mean ± SD, *n* = 3) represent the specific binding of UNC0642.

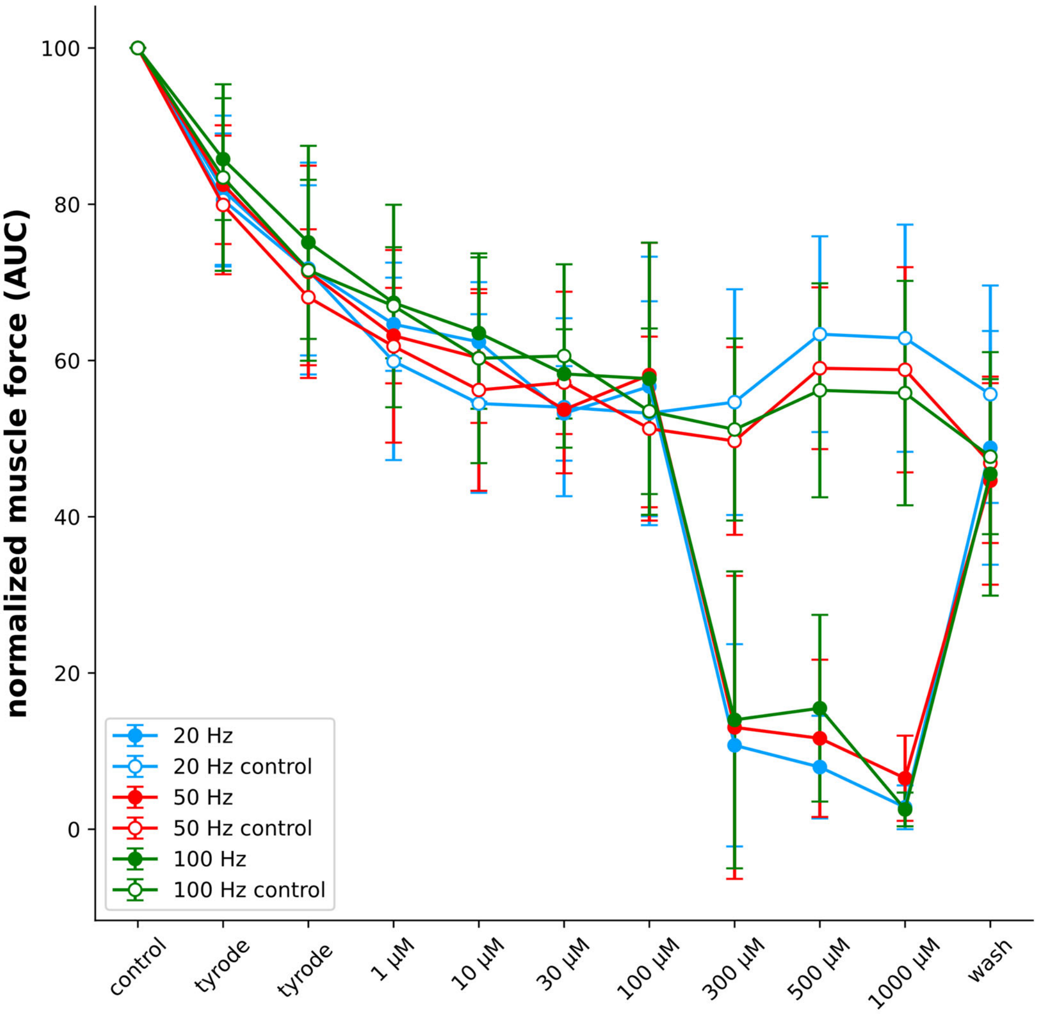


**Figure S8:** Normalized rat muscle force in the presence of increasing cycloguanil (**6**) concentrations and in the absence of cycloguanil (**6**) (control) during indirect electric field stimulations of 20, 50, and 100 Hz. Error bars indicate the standard deviation (*n* is between 3 and 13).

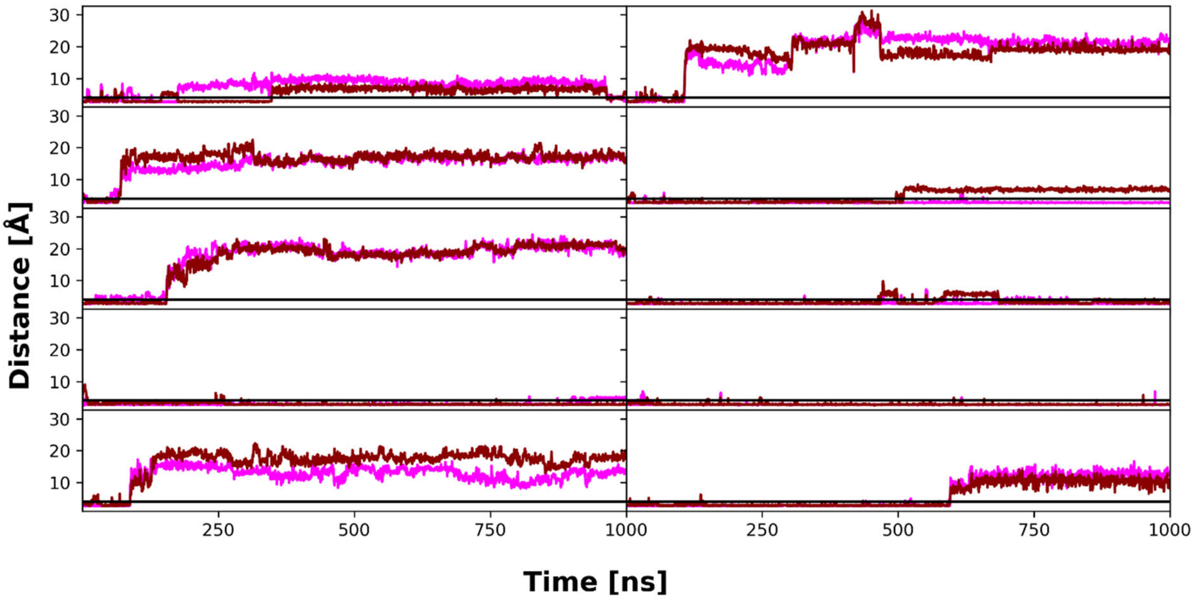


**Figure S9:** Distance of nitrogens that can act as hydrogen bond donors of cycloguanil to side chain oxygens of E62$_\gamma$ (pink) and E200$_\gamma$ (dark red) during 10 replicas of unbiased MD simulations.

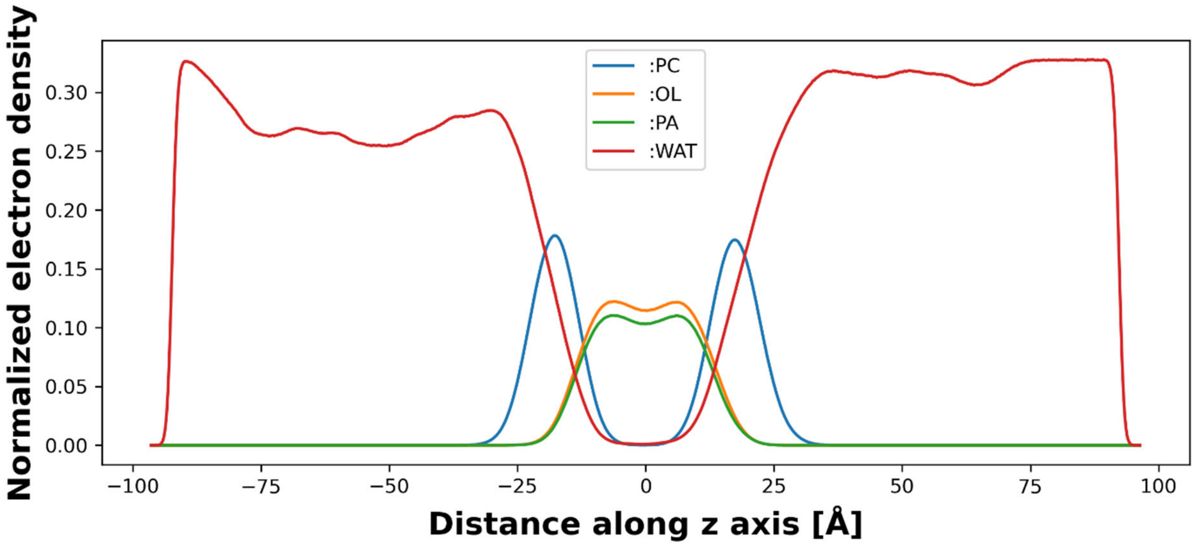


**Figure S10:** Normalized (number of electrons per volume [$e^-$/Å$^3$]) electron density of membrane components and water averaged over all 10 replicas of 1 µs long MD simulations. The electron density is plotted against the z-coordinate, which is parallel to the membrane normal. The membrane is centered at 0 Å for phosphatidylcholine (:PC), oleic acid (:OL), palmitoyl acid (:PA), and water (:WAT).

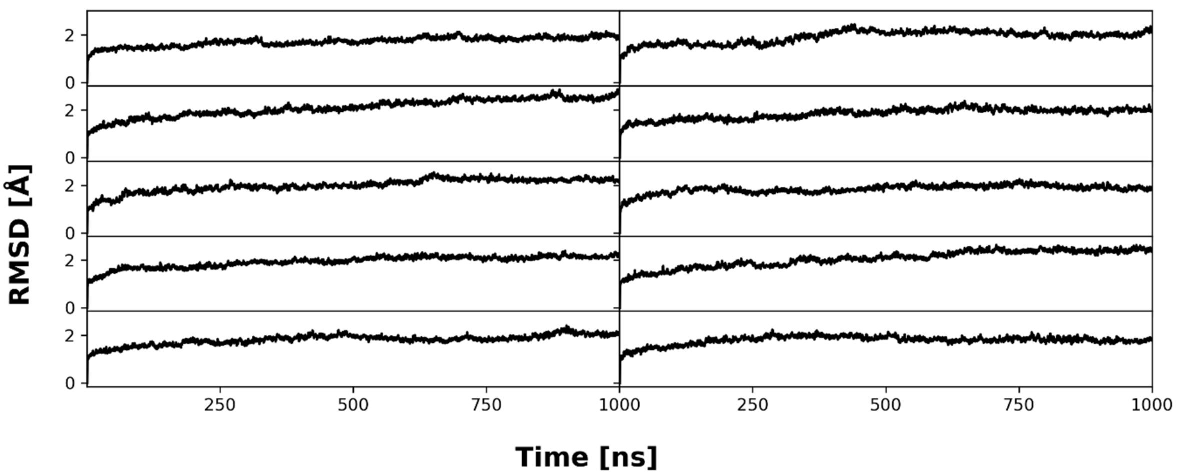


**Figure S11:** Backbone (C, CA, N) RMSD of nAChR during 10 replicas of 1 µs long MD simulations with respect to the first frame of the production run.

### Supplemental Tables

**Table S1:** List of commercially obtained test compounds.

| # | Compound | Salt | Purity | Supplier |
|---|---|---|---|---|
| 1 | Bunazosin | - | ≥ 95% | Angene (Honk Kong, China) |
| 2 | C-021 | - | ≥ 97% | BIONET – KeyOrganics Ltd (Cornwall, United Kingdom) |
| 3 | Cycloguanil | HCl | ≥ 95% | Otava Ltd (Kiew, Ukraine) |
| 4 | MS012 | - | ≥ 90% | TimTec, LLC (Newark, United States) |
| 5 | PTMD01-0019C | - | ≥ 85% | ChemBridge Corporation (San Diego, United States) |
| 6 | PTMD01-0020C | - | ≥ 90% | Enamine Ltd (Kiew, Ukraine) |
| 7 | PTMD01-0021C | - | ≥ 85% | ChemBridge Corporation (San Diego, United States) |
| 8 | PTMD01-0022C | - | ≥ 85% | ChemBridge Corporation (San Diego, United States) |
| 9 | PTMD01-0023C | - | ≥ 90% | Enamine Ltd (Kiew, Ukraine) |
| 10 | PTMD01-0024C | - | ≥ 85% | ChemBridge Corporation (San Diego, United States) |
| 11 | PTMD01-0025C | - | ≥ 90% | Enamine Ltd (Kiew, Ukraine) |
| 12 | PTMD99-0001C | HI | ≥ 90% | Enamine Ltd (Kiew, Ukraine) |
| 13 | PTMD99-0002C | - | ≥ 90% | UkrOrgSynthesis Ltd. (Kiew, Ukraine) |
| 14 | PTMD99-0004C | 2 HCl | ≥ 85% | ChemBridge Corporation (San Diego, United States) |
| 15 | PTMD99-0005C | - | ≥ 90% | ChemBridge Corporation (San Diego, United States) |
| 16 | PTMD99-0006C | - | ≥ 85% | ChemBridge Corporation (San Diego, United States) |
| 17 | PTMD99-0008C | $H_2SO_4$ | ≥ 90% | Vitas-M Laboratory Ltd (Causeway Bay, Hong Kong) |
| 18 | PTMD99-0009C | HBr | ≥ 90% | Enamine Ltd (Kiew, Ukraine) |
| 19 | PTMD99-0010C | 2 HCl | ≥ 85% | ChemBridge Corporation (San Diego, United States) |
| 20 | PTMD99-0011C | - | ≥ 85% | ChemBridge Corporation (San Diego, United States) |
| 21 | PTMD99-0013C | HCl | ≥ 90% | ChemBridge Corporation (San Diego, United States) |
| 22 | PTMD99-0014C | HCl | ≥ 90% | Enamine Ltd (Kiew, Ukraine) |
| 23 | PTMD99-0015C | - | ≥ 85% | ChemBridge Corporation (San Diego, United States) |
| 24 | PTMD99-0016C | - | ≥ 85% | ChemBridge Corporation (San Diego, United States) |
| 25 | PTMD99-0020C | - | ≥ 85% | ChemBridge Corporation (San Diego, United States) |
| 26 | PTMD99-0021C | - | ≥ 85% | ChemBridge Corporation (San Diego, United States) |
| 27 | PTMD99-0023C | 2 HCl | ≥ 85% | ChemBridge Corporation (San Diego, United States) |
| 28 | PTMD99-0024C | 2 HCl | ≥ 85% | ChemBridge Corporation (San Diego, United States) |
| 29 | PTMD99-0025C | 2 HCl | ≥ 85% | ChemBridge Corporation (San Diego, United States) |
| 30 | PTMD99-0026C | 2 HCl | ≥ 85% | ChemBridge Corporation (San Diego, United States) |
| 31 | PTMD99-0028C | 2 HCl | ≥ 85% | ChemBridge Corporation (San Diego, United States) |
| 32 | PTMD99-0029C | 2 HCl | ≥ 85% | ChemBridge Corporation (San Diego, United States) |
| 33 | PTMD99-0031C | 2 HCl | ≥ 85% | ChemBridge Corporation (San Diego, United States) |
| 34 | PTMD99-0032C | HCl | ≥ 90% | Maybridge Ltd (Cornwall, United Kingdom) |
| 35 | PTMD99-0035C | HCl | ≥ 90% | Otava Ltd (Kiew, Ukraine) |
| 36 | PTMD99-0036C | HCl | ≥ 90% | ChemBridge Corporation (San Diego, United States) |
| 37 | PTMD99-0038C | HCl | ≥ 95% | Otava Ltd (Kiew, Ukraine) |
| 38 | PTMD99-0041C | HCl | ≥ 85% | ChemBridge Corporation (San Diego, United States) |
| 39 | PTMD99-0044C | HCl | ≥ 90% | Maybridge Ltd (Cornwall, United Kingdom) |
| 40 | PTMD99-0045C | HCl | ≥ 90% | Maybridge Ltd (Cornwall, United Kingdom) |
| 41 | UNC0379 | - | ≥ 98% | TargetMol (Boston, United States) |
| 42 | ZT-12-037-01 | - | ≥ 98% | TargetMol (Boston, United States) |

**Table S2:** All analogs of UNC0646 tested for affinity to MB327-PAM-1 in nAChR determined in MS Binding Assays based on a two-dimensional similarity search.

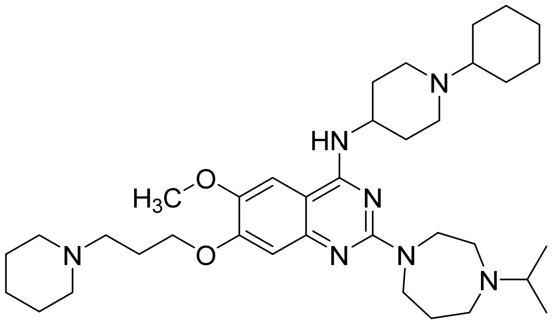

**UNC0646**

**$pK_i$ = 5.83 ± 0.05**[a]

**Remaining reporter ligand binding: 21 ± 3 %**[b]

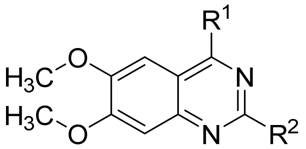

| Compound | $R^1$ | $R^2$ | Remaining reporter ligand binding [%][b] |
|---|---|---|---|
| C-021 | 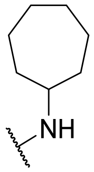 | 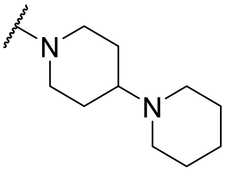 | 63 ± 2 |
| ZT-12-037-01 | 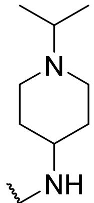 | 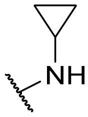 | 66 ± 5 |
| MS012 | 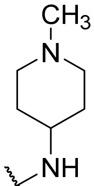 | 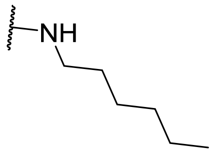 | 83 ± 6 |
| UNC0379 | 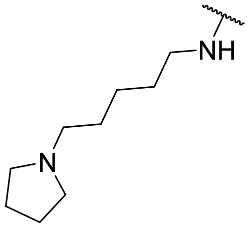 | 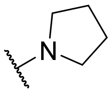 | 59 ± 4 |
| Bunazosin | 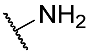 | 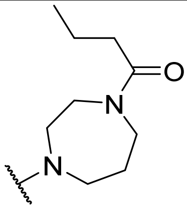 | 90 ± 7 |

| | | | |
|---|---|---|---|
| PTMD01-0019C | 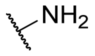 | 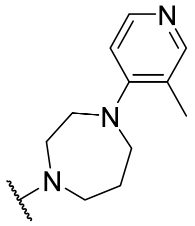 | 47 ± 3 |
| PTMD01-0020C | 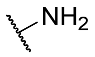 | 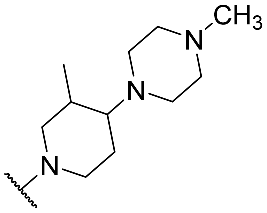 | 73 ± 7 |
| PTMD01-0021C | 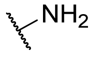 | 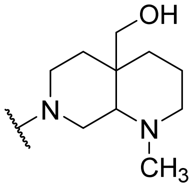 | 87 ± 2 |
| PTMD01-0022C | 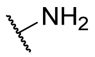 | 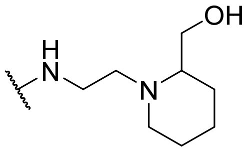 | 108 ± 10 |
| PTMD01-0023C | 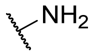 | 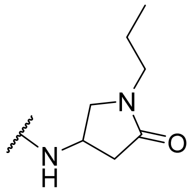 | 98 ± 3 |
| PTMD01-0024C | 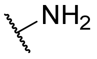 | 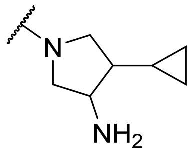 | 83 ± 4 |
| PTMD01-0025C | 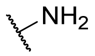 | 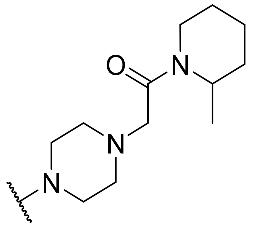 | 81 ± 5 |

[a] The $pK_i$ value of UNC0646 has been reported in ref. [5].

[b] Characterized by UNC0642 MS Binding Assays; Percentage of remaining reporter ligand binding in the presence of test compounds as compared to 100% reporter ligand binding in the absence of a competitor. Results are based on thirty measurements for UNC0646 and three measurements for all other compounds at a test compound concentration of 10 µM and a reporter ligand concentration of 1 µM. Mean and standard deviation are displayed.

**Table S3:** Results of affinity testing to MB327-PAM-1 in nAChR determined in MS Binding Assays of initial compounds ordered based on structure-based screening.

| Name | Structure | Remaining reporter ligand binding [%][a] |
|---|---|---|
| PTMD99-0001C (**3**) | 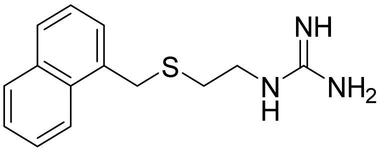 | 88 ± 5 |
| PTMD99-0002C | 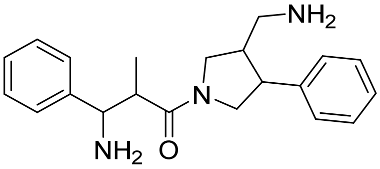 | 101 ± 9 |
| PTMD99-0004C | 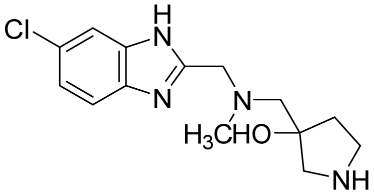 | 92 ±7 |
| PTMD99-0005C | 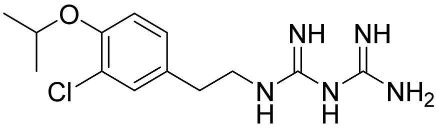 | 101 ± 3 |
| PTMD99-0006C (**13**) | 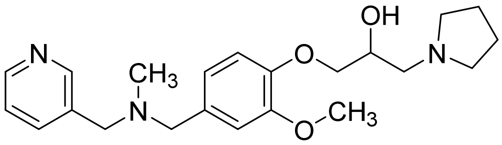 | 93 ± 7 |
| PTMD99-0008C | 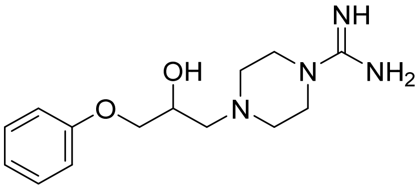 | 95 ± 4 |
| PTMD99-0009C | 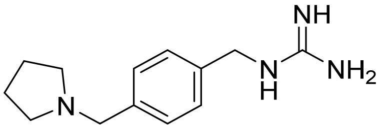 | 92 ± 9 |
| PTMD99-0010C (**14**) | 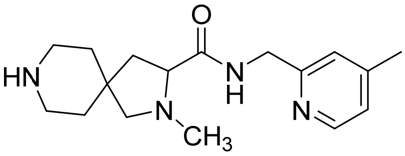 | 96 ± 5 |
| PTMD99-0011C | 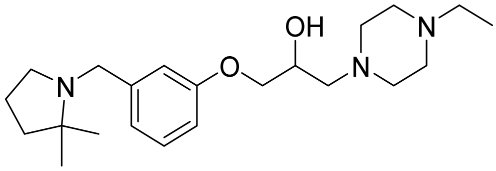 | 95 ± 7 |
| PTMD99-0013C | 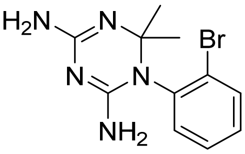 | 101 ± 6 |

| | | |
|---|---|---|
| PTMD99-0014C **(15)** | 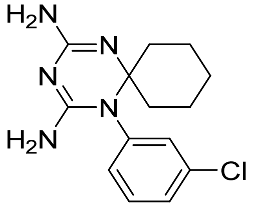 | 94 ± 6 |
| PTMD99-0015C | 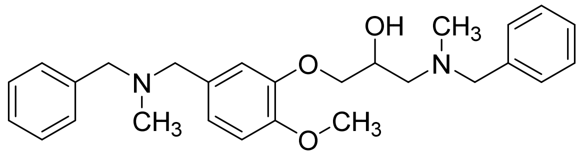 | 98 ± 8 |

[a] Characterized by UNC0642 MS Binding Assays; Percentage of remaining reporter ligand binding in the presence of test compounds as compared to 100% reporter ligand binding in the absence of a competitor. If not stated otherwise, results are based on three measurements at a test compound concentration of 10 µM and a reporter ligand concentration of 1 µM. Mean and standard deviation are displayed.

**Table S4:** Results of affinity testing of compounds analogous to the best hits from the initially ordered compounds based on structure-based screening.

| Name | Structure | Remaining reporter ligand binding [%][a] |
|---|---|---|
| PTMD99-0016C (**4**) | 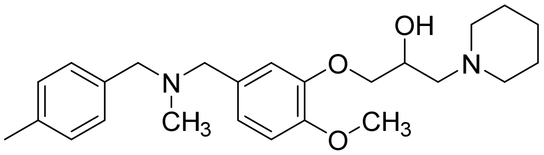 | 78 ± 3 |
| PTMD99-0020C (**16**) | 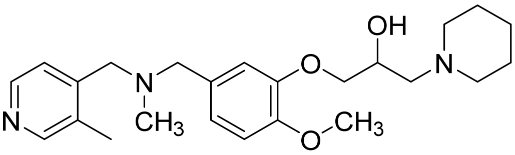 | 97 ± 4 |
| PTMD99-0021C | 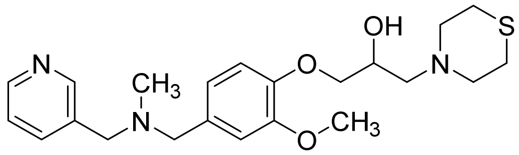 | 107 ± 11 |
| PTMD99-0023C (**17**) | 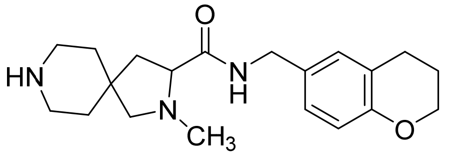 | 108 ± 4 |
| PTMD99-0024C (**18**) | 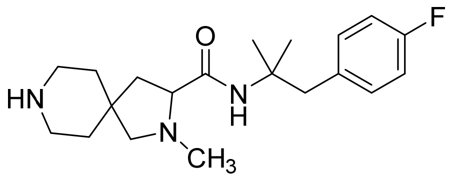 | 99 ± 6 |
| PTMD99-0025C (**19**) | 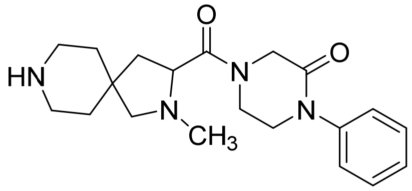 | 100 ± 6 |
| PTMD99-0026C (**5**) | 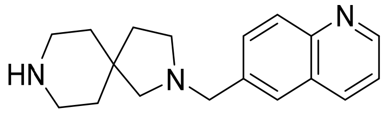 | 90 ± 5 |
| PTMD99-0028C (**20**) | 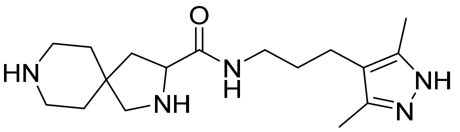 | 101 ± 7 |
| PTMD99-0029C | 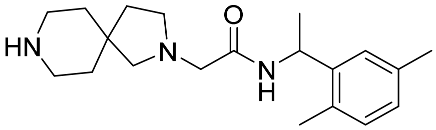 | 98 ± 4 |
| PTMD99-0031C (**21**) | 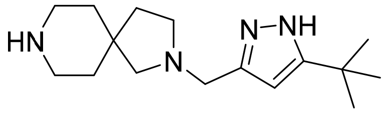 | 101 ± 8 |
| PTMD99-0032C (**22**) | 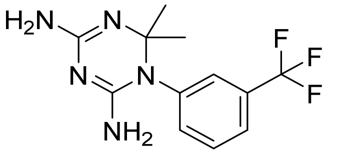 | 98 ± 6 |

| | | |
|---|---|---|
| PTMD99-0035C (**23**) | 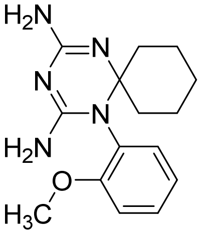 | 109 ± 8 |
| PTMD99-0036C (**24**) | 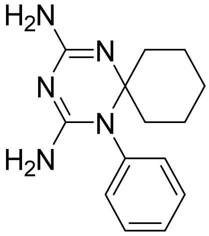 | 93 ± 4 |
| PTMD99-0038C (**25**) | 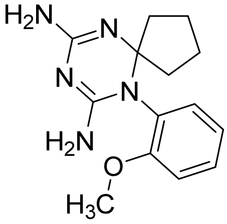 | 100 ± 4 |
| Cycloguanil (**6**) | 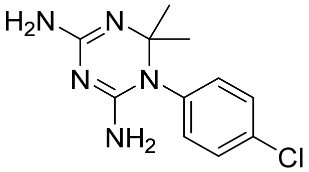 | 76 ± 3 |
| PTMD99-0041C (**26**) | 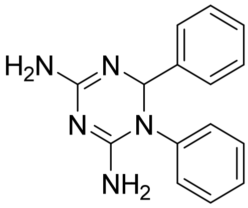 | 81 ± 3 |
| PTMD99-0044C (**27**) | 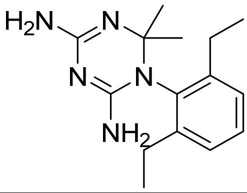 | 106 ± 6 |
| PTMD99-0045C (**28**) | 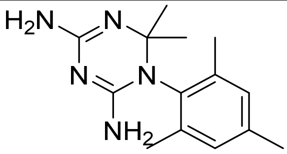 | 95 ± 8 |

[a] Characterized by UNC0642 MS Binding Assays; Percentage of remaining reporter ligand binding in the presence of test compounds as compared to 100% reporter ligand binding in the absence of a competitor. If not stated otherwise, results are based on three measurements at a test compound concentration of 10 µM and a reporter ligand concentration of 1 µM. Mean and standard deviation are displayed.

**Table S5:** Restoration of muscle force in soman-inhibited rat muscles after cyclouanil (**6**) treatment.[a]

| | 20 Hz | | | 50 Hz | | | 100 Hz | | |
|---|---|---|---|---|---|---|---|---|---|
| | Mean [%] | SD [%] | n | Mean [%] | SD [%] | n | Mean [%] | SD [%] | n |
| control | 100.00 | 0.00 | 27 | 100.00 | 0.00 | 27 | 100.00 | 0.00 | 27 |
| soman | 4.30 | 6.02 | 27 | 3.86 | 8.36 | 27 | 4.51 | 8.95 | 27 |
| wash | 0.00 | 0.00 | 27 | 0.00 | 0.00 | 27 | 0.00 | 0.00 | 27 |
| 1 µM | 1.86 | 2.91 | 13 | 0.64 | 1.29 | 13 | 0.40 | 1.00 | 13 |
| 10 µM | 6.20 | 8.40 | 18 | 0.82 | 1.28 | 18 | 0.25 | 0.56 | 18 |
| 30 µM | 7.66 | 7.64 | 5 | 1.38 | 2.34 | 5 | 1.14 | 2.51 | 5 |
| 70 µM | 30.87 | 19.23 | 5 | 2.76 | 3.51 | 5 | 0.68 | 1.32 | 5 |
| 100 µM | 30.42 | 18.04 | 27 | 4.68 | 4.76 | 27 | 1.29 | 2.72 | 27 |
| 150 µM | 7.78 | 10.79 | 9 | 0.85 | 1.33 | 9 | 0.28 | 0.57 | 9 |
| 200 µM | 1.69 | 3.35 | 4 | 0.23 | 0.41 | 4 | 0.00 | 0.00 | 4 |
| 300 µM | 0.51 | 2.09 | 17 | 0.51 | 1.69 | 17 | 0.71 | 2.44 | 17 |
| 500 µM | 0.24 | 0.53 | 5 | 1.37 | 3.05 | 5 | 1.67 | 3.73 | 5 |
| 1000 µM | 0.00 | 0.00 | 5 | 0.73 | 1.62 | 5 | 1.05 | 2.35 | 5 |
| wash | 14.30 | 15.66 | 27 | 2.43 | 3.89 | 27 | 0.39 | 1.44 | 27 |

[a] For the stimulation frequencies of 20, 50, and 100 Hz, the mean muscle force restoration, the standard deviation (SD), and the number of experiments (*n*) are displayed at each testing point.

### Analytical Data

**6-Methoxy-2-(piperidin-1-yl)-7-[3-(piperidin-1-yl)propoxy]-*N*-[5-(pyrrolidin-1-yl)pentyl]quinazolin-4-amine (1k):** mp.: 54 °C. $R_f$ = 0.34 [10% 3 M $NH_3$ (in MeOH) in $CH_2Cl_2$]. IR (film): ṽ = 2935, 1581, 1493, 1244, 754 $cm^{-1}$. $^1H$ NMR (500 MHz, $CD_2Cl_2$): δ = 1.37-1.47 (m, 2 H, $CH_2CH_2CH_2NCH_2CH_2CH_2O$), 1.48-1.55 (m, 2 H, $CH_2CH_2CH_2NH$), 1.55-1.61 (m, 8 H, $CH_2CH_2CH_2NC$, $CH_2CH_2CH_2NCH_2CH_2CH_2O$), 1.62-1.70 (m, 4 H, $CH_2CH_2CH_2NC$, $CH_2CH_2CH_2CH_2NH$), 1.74 (p, *J* = 7.0 Hz, 2 H, $CH_2CH_2NH$), 1.80-1.89 [m, 4 H, $CH_2CH_2N(CH_2)_5NH$], 2.00 (p, *J* = 6.9 Hz, 2 H, $CH_2CH_2O$), 2.22-2.42 (m, 4 H, $CH_2NCH_2CH_2CH_2O$), 2.42-2.48 (m, 2 H, $CH_2CH_2CH_2O$), 2.57-2.66 [m, 2 H, $CH_2(CH_2)_4NH$], 2.66-2.84 [m, 4 H, $CH_2N(CH_2)_5NH$], 3.53-3.64 (m, 2 H, $CH_2NH$), 3.73-3.84 (m, 4 H, $CH_2NC$), 3.89 (s, 3 H, $CH_3O$), 4.11 (t, *J* = 6.7 Hz, 2 H, $CH_2O$), 5.77 (s, 1 H, NH), 6.84 (s, 1 H, CHCOCH$_2$), 6.98 (s, 1 H, CHCOCH$_3$). $^{13}C$ NMR (126 MHz, $CD_2Cl_2$): δ = 23.82 [$CH_2CH_2N(CH_2)_5NH$], 24.94 ($CH_2CH_2CH_2NCH_2CH_2CH_2O$), 25.17 ($CH_2CH_2CH_2NH$), 25.57 ($CH_2CH_2CH_2NC$), 26.41 ($CH_2CH_2NCH_2CH_2CH_2O$)*, 26.46 ($CH_2CH_2NC$)*, 27.05 ($CH_2CH_2O$), 28.02 [$CH_2(CH_2)_3NH$], 29.19 ($CH_2CH_2NH$), 41.36 ($CH_2NH$), 45.39 ($CH_2NC$), 54.41 [$CH_2N(CH_2)_5NH$], 55.00 ($CH_2NCH_2CH_2CH_2O$), 55.97 ($CH_2CH_2CH_2O$), 56.31 [$CH_2(CH_2)_4NH$], 56.96 ($CH_3O$), 67.57 ($CH_2O$), 102.41 (CHCOCH$_3$), 103.30 (CCNH), 106.82 (CHCOCH$_2$), 145.83 (COCH$_3$), 149.62 (CCCNH), 154.35 (COCH$_2$), 159.04 (CNCNH), 159.49 (CNH). HRMS-EI m/z $[M]^+$ calcd. for $C_{31}H_{50}N_6O_2$: 538.3995, found: 538.3989.

***N*$^1$-Cyclohexyl-*N*$^2$-{6-methoxy-7-[3-(piperidin-1-yl)propoxy]quinazolin-4-yl}-*N*$^1$-methylethane-1,2-diamine (2b):** $R_f$ = 0.38 [10% 3 M $NH_3$ (in MeOH) in $CH_2Cl_2$]. IR (film): ṽ = 3020, 1641, 1215 $cm^{-1}$. $^1H$ NMR (500 MHz, $CD_2Cl_2$): δ = 1.03-1.16 (m, 1 H, $CH_2CH_2CH_2CHN$), 1.19-1.37 (m, 4 H, $CH_2CH_2CHN$), 1.37-1.47 (m, 2 H, $CH_2CH_2CH_2NCH_2CH_2CH_2O$), 1.47-1.59 (m, 4 H, $CH_2CH_2NCH_2CH_2CH_2O$), 1.59-1.66 (m, 1 H, $CH_2CH_2CH_2CHN$), 1.77-1.87 (m, 4 H, $CH_2CH_2CHN$), 2.02 (p, *J* = 6.8 Hz, 2 H, $CH_2CH_2O$), 2.32 (s, 3 H, $CH_3N$), 2.34-2.43 (m, 4 H, $CH_2NCH_2CH_2CH_2O$), 2.43-2.52 (m, 3 H, $CH_2CH_2CH_2O$, $CH_2CHN$), 2.68-2.88 (m, 2 H, $CH_2CH_2NH$), 3.53-3.67 (m, 2 H, $CH_2NH$), 3.94 (s, 3 H, $CH_3O$), 4.16 (t, *J* = 6.7 Hz, 2 H, $CH_2O$), 6.50 (d, *J* = 13.6 Hz, 1 H, NH), 6.95 (s, 1 H, CHCOCH$_3$), 7.14 (s, 1 H, CHCOCH$_2$), 8.43 (s, 1 H, CHNCNH). $^{13}C$ NMR (126 MHz, $CD_2Cl_2$): δ = 24.97 ($CH_2CH_2CH_2NCH_2CH_2CH_2O$), 26.35 ($CH_2CH_2CHN$), 26.52 ($CH_2CH_2NCH_2CH_2CH_2O$), 26.68 ($CH_2CH_2O$), 27.02 ($CH_2CH_2CH_2CHN$), 29.26 ($CH_2CHN$), 37.08 ($CH_3N$), 38.38 ($CH_2NH$), 52.08 ($CH_2CH_2NH$), 55.03 ($CH_2NCH_2CH_2CH_2O$), 55.91 ($CH_2CH_2CH_2O$), 56.39 ($CH_3O$), 63.59 ($CH_2CHN$), 67.87 ($CH_2O$), 100.42 (CHCOCH$_3$), 108.83 (CHCOCH$_2$), 109.02 (14), 146.84 (15), 149.46 (12), 154.21 (11), 154.49 (CHNCNH), 158.65 (19). HRMS-ESI m/z $[M+H]^+$ calcd. for $C_{26}H_{42}N_5O_2$: 456.3339, found: 456.3334.

**6-Methoxy-*N*-[3-(4-methylpiperazin-1-yl)butyl]-7-[3-(piperidin-1-yl)propoxy]quinazolin-4-amine (2c):** mp.: 85 °C. $R_f$ = 0.22 [10% 3 M $NH_3$ (in MeOH) in $CH_2Cl_2$]. IR (film): ṽ = 2100, 1622, 1504, 1338, 1217 $cm^{-1}$. $^1H$ NMR (500 MHz, $CD_2Cl_2$): δ = 1.05 (d, *J* = 6.6 Hz, 3 H, $CH_3CH$), 1.37-1.49 (m, 2 H, $CH_2CH_2CH_2NCH_2CH_2CH_2O$), 1.51-1.61 (m, 4 H, $CH_2CH_2NCH_2CH_2CH_2O$), 1.65-1.97 (m, 2 H, $CH_2CH_2NH$), 1.98-2.08 (m, 2 H, $CH_2CH_2O$), 2.26 (s, 3 H, $CH_3N$), 2.28-2.62 (m, 12 H, $CH_2CH_2NCH_3$, $CH_2NCH_2CH_2CH_2O$), 2.64-2.77 (m, 2 H, $CH_2CH_2NCH_3$), 2.78-2.96 (m, 1 H, $CHCH_3$), 3.58-3.70 (m, 1 H, $CH_2NH$), 3.72-3.85 (m, 1 H, $CH_2NH$), 3.97 (s, 3 H, $CH_3O$), 4.17 (t, *J* = 6.6 Hz, 2 H, $CH_2O$), 6.71 (s, 1 H, NH), 7.05 (s, 1 H, CHCOCH$_3$), 7.15 (s, 1 H, CHCOCH$_2$), 8.43 (s, 1 H, CHNCNH). $^{13}C$ NMR (126 MHz, $CD_2Cl_2$): δ = 13.74 ($CH_3CH$), 24.92 ($CH_2CH_2CH_2NCH_2CH_2CH_2O$), 26.47 ($CH_2CH_2NCH_2CH_2CH_2O$), 26.98 ($CH_2CH_2O$), 32.20 ($CH_2CH_2NH$), 40.71 ($CH_2NH$), 46.07 ($CH_3N$), 48.75 ($CH_2CH_2NCH_3$), 55.01 ($CH_2NCH_2CH_2CH_2O$), 55.93 ($CH_2NCH_3$, $CH_2CH_2CH_2O$), 57.61 ($CH_3O$), 59.32 ($CHCH_3$), 67.81 ($CH_2O$), 102.33 (CHCOCH$_3$), 108.99 (CHCOCH$_2$), 109.06 (CCNH), 147.20 (CCCNH), 149.40 (COCH$_3$), 154.51 (CHNCNH, COCH$_2$), 158.94 (CNH). HRMS-ESI m/z $[M+H]^+$ calcd. for $C_{26}H_{43}N_6O_2$: 471.3447, found: 471.3443.

**6-Methoxy-7-[3-(piperidin-1-yl)propoxy]-4-[4-(pyrrolidin-1-yl)piperidin-1-yl]quinazoline (2d):** IR (film): ṽ = 2937, 1643, 1504, 1209 cm$^{-1}$. $^1$H NMR (500 MHz, $CD_2Cl_2$): δ = 1.36-1.49 (m, 2 H, $CH_2CH_2CH_2NCH_2CH_2CH_2O$), 1.49-1.59 (m, 4 H, $CH_2CH_2NCH_2CH_2CH_2O$), 1.69-1.82 (m, 6 H, $CH_2CHN$, $CH_2CH_2NCH$), 1.98-2.10 (m, 4 H, $CH_2CHN$, $CH_2CH_2O$), 2.28 (tt, *J* = 10.2, 4.0 Hz, 1 H, $CHNCH_2$), 2.32-2.43 (m, 4 H, $CH_2NCH_2CH_2CH_2O$), 2.46 (t, *J* = 7.1 Hz, 2 H, $CH_2CH_2CH_2O$), 2.51-2.64 (m, 4 H, $CH_2NCH$), 3.04-3.15 (m, 2 H, $CH_2NC$), 3.94 (s, 3 H, $CH_3O$), 4.04-4.14 (m, 2 H, $CH_2NC$), 4.18 (t, *J* = 6.7 Hz, 2 H, $CH_2O$), 7.11 (s, 1 H, $CHCOCH_3$), 7.19 (s, 1 H, $CHCOCH_2$), 8.55 (s, 1 H, CHNCN). $^{13}$C NMR (126 MHz, $CD_2Cl_2$): δ = 23.74 ($CH_2CH_2NCH$), 24.96 ($CH_2CH_2CH_2NCH_2CH_2CH_2O$), 26.52 ($CH_2CH_2NCH_2CH_2CH_2O$), 26.98 ($CH_2CH_2O$), 32.07 ($CH_2CHN$), 49.09 ($CH_2CH_2CHN$), 51.78 ($CH_2NCH$), 55.03 ($CH_2NCH_2CH_2CH_2O$), 55.89 ($CH_2CH_2CH_2O$), 56.31 ($CH_3O$), 62.21 ($CHNCH_2$), 67.92 ($CH_2O$), 103.89 ($CHCOCH_3$), 108.49 ($CHCOCH_2$), 111.84 ($CCHCOCH_3$), 149.03 ($COCH_3$), 149.47 ($CCCNCH_2$), 153.34 (CHNCN), 154.41 ($COCH_2$), 164.49 ($CNCH_2$). HRMS-ESI m/z [M+H]$^+$ calcd. for $C_{26}H_{40}N_5O_2$: 454.3182, found: 454.3177.

***N*-[1-(Azepan-1-yl)-2-methylpropan-2-yl]-6-methoxy-7-[3-(piperidin-1-yl)propoxy]quinazolin-4-amine (2e):** Hygroscopic. $R_f$ = 0.30 [5% 7 M $NH_3$ (in MeOH) in $CH_2Cl_2$]. IR (film): ṽ = 2926, 1643, 1498 cm$^{-1}$. $^1$H NMR (500 MHz, $CD_2Cl_2$): δ = 1.38-1.47 (m, 2 H, $CH_2CH_2CH_2NCH_2CH_2CH_2O$), 1.52-1.62 (m, 10 H, $CH_3C$, $CH_2CH_2NCH_2CH_2CH_2O$), 1.62-1.78 (m, 8 H, $CH_2CH_2CH_2NCH_2CNH$), 2.03 (p, *J* = 6.9 Hz, 2 H, $CH_2CH_2O$), 2.26-2.46 (m, 4 H, $CH_2NCH_2CH_2CH_2O$), 2.46-2.57 (m, 2 H, $CH_2CH_2CH_2O$), 2.75 (s, 2 H, $CH_2CNH$), 2.82-2.96 (m, 4 H, $CH_2NCH_2CNH$), 3.95 (s, 3 H, $CH_3O$), 4.16 (t, *J* = 6.6 Hz, 2 H, $CH_2O$), 6.84 (s, 1 H, NH), 6.93 (s, 1 H, $CHCOCH_3$), 7.12 (s, 1 H, $CHCOCH_2$), 8.40 (s, 1 H, CHNCNH). $^{13}$C NMR (126 MHz, $CD_2Cl_2$): δ = 24.91 ($CH_2CH_2CH_2NCH_2CH_2CH_2O$), 25.77 ($CH_3C$), 26.45 ($CH_2CH_2NCH_2CH_2CH_2O$), 26.97 ($CH_2CH_2O$), 27.51 ($CH_2CH_2CH_2NCH_2CNH$), 29.59 ($CH_2CH_2NCH_2CNH$), 54.36 ($CH_3C$), 55.00 ($CH_2NCH_2CH_2CH_2O$), 55.91 ($CH_2CH_2CH_2O$), 56.38 ($CH_3O$), 58.82 ($CH_2NCH_2CNH$), 67.81 ($CH_2O$), 69.97 ($CH_2CNH$), 100.45 ($CHCOCH_3$), 109.02 ($CHCOCH_2$), 109.86 (CCNH), 146.93 (CCCNH), 149.32 ($COCH_3$), 153.88 ($COCH_2$), 154.04 (CHNCNH), 158.64 (NCNH). HRMS-ESI m/z [M+H]$^+$ calcd. for $C_{27}H_{44}N_5O_2$: 470.3495, found: 470.3492.

***N*-(1-Propan-2-ylpiperidin-4-yl)-6-methoxy-7-[3-(piperidin-1-yl)propoxy]quinazolin-4-amine (2f):** mp.: 166 °C. $R_f$ = 0.18 [10% 4 M $NH_3$ (in MeOH) in $CH_2Cl_2$]. IR (film): ṽ = 2933, 1593, 1506, 1254 cm$^{-1}$. $^1$H NMR (500 MHz, $CD_3OD$): δ = 1.14 (d, *J* = 6.6 Hz, 6 H, $CH_3CH$), 1.45-1.56 (m, 2 H, $CH_2CH_2CH_2NCH_2CH_2CH_2O$), 1.57-1.69 (m, 4 H, $CH_2CH_2NCH_2CH_2CH_2O$), 1.69-1.81 (m, 2 H, $CH_2CHNH$), 2.04-2.18 (m, 4 H, $CH_2CH_2O$, $CH_2CHNH$), 2.38-2.48 (m, 2 H, $CH_2NCH$), 2.48-2.60 (m, 4 H, $CH_2NCH_2CH_2CH_2O$), 2.60-2.69 (m, 2 H, $CH_2CH_2CH_2O$), 2.82 (hept, *J* = 6.5 Hz, 1 H, $CH_3CH$), 2.98-3.09 (m, 2 H, $CH_2NCH$), 3.97 (s, 3 H, $CH_3O$), 4.09-4.30 (m, 3 H, $CH_2O$, CHNH), 7.07 (s, 1 H, CHCN), 7.60 (s, 1 H, CHCCN), 8.30 (s, 1 H, CHNCNH). $^{13}$C NMR (126 MHz, $CD_3OD$): δ = 18.47 ($CH_3CH$), 25.04 ($CH_2CH_2CH_2NCH_2CH_2CH_2O$), 26.38 ($CH_2CH_2NCH_2CH_2CH_2O$), 27.00 ($CH_2CH_2O$), 32.37 ($CH_2CHNH$), 48.83 ($CH_2NCH$), 49.84 (CHNH), 55.46 ($CH_2NCH_2CH_2CH_2O$), 56.06 ($CH_3CH$), 56.82 ($CH_3O$), 57.00 ($CH_2CH_2CH_2O$), 68.32 ($CH_2O$), 102.94 (CHCCN), 107.81 (CHCN), 110.17 (CCNH), 146.65 (CCCNH), 150.87 ($CH_3OC$), 154.28 (CHNCNH), 155.48 ($CH_2OC$), 159.65 (CNH). HRMS-ESI m/z [M+H]$^+$ calcd. for $C_{25}H_{40}N_5O_2$: 442.3182, found: 442.3175.

***N*-(1-Propan-2-ylpiperidin-4-yl)-6-methoxy-7-[3-(piperidin-1-ylmethyl)pyrrolidin-1-yl]quinazolin-4-amine (2g):** mp.: 244 °C. $R_f$ = 0.35 [10% 3 M $NH_3$ (in MeOH) in $CH_2Cl_2$]. IR (film): ṽ = 2935, 1612, 1504, 1363 cm$^{-1}$. $^1$H NMR (500 MHz, $CD_2Cl_2$): δ = 1.04 (d, *J* = 6.6 Hz, 6 H, $CH_3CH$), 1.33-1.47 (m, 2 H, $CH_2CH_2CH_2N$), 1.47-1.60 (m, 6 H, $CH_2CH_2CH_2N$, $CH_2CHNH$), 1.60-1.78 (m, 1 H, $CH_2CH_2CHCH_2N$), 2.02-2.09 (m, 1 H, $CH_2CH_2CHCH_2N$), 2.09-2.17 (m, 2 H, $CH_2CHNH$), 2.19-2.44 (m, 8 H, $CH_2NCH_2CHCH_2N$,

$CH_2CH_2CHNH$), 2.49 (hept, *J* = 7.0 Hz, 1 H, $CHCH_2N$), 2.76 (hept, *J* = 6.4 Hz, 1 H, $CH_3CH$), 2.83-2.93 (m, 2 H, $CH_2CH_2CHNH$), 3.26 (dd, *J* = 10.2, 7.2 Hz, 1 H, $CHCH_2NC$), 3.51-3.62 (m, 3 H, $CH_2NC$), 3.90 (s, 3 H, $CH_3O$), 4.10-4.26 (m, 1 H, CHNH), 5.01-5.18 (m, 1 H, NH), 6.76 (s, 1 H, $CHCOCH_3$), 6.79 (s, 1 H, $CHCNCH_2$), 8.34 (s, 1 H, CHNC). $^{13}$C NMR (126 MHz, $CD_2Cl_2$): $\delta$ = 18.46 ($CH_3CH$), 24.97 ($CH_2CH_2CH_2N$), 26.55 ($CH_2CH_2CH_2N$), 30.45 ($CH_2CHCH_2N$), 33.37 ($CH_2CHNH$), 36.58 ($CHCH_2N$), 48.05 ($CH_2CH_2CHNH$), 48.85 (CHNH), 50.56 ($CH_2CH_2CHCH_2N$), 54.85 ($CH_3CH$), 55.41 ($CH_2CH_2CH_2N$), 55.84 ($CNCH_2CH$), 56.33 ($CH_3O$), 63.20 ($CH_2CHCH_2NC$), 100.12 ($CHCOCH_3$), 106.53 (CCNH), 109.46 ($CHCNCH_2$), 145.49 ($CNCH_2$), 147.48 (CCCNH), 150.16 ($COCH_3$), 154.18 (CHNCNH), 157.71 (CNH). HRMS-ESI m/z $[M+H]^+$ calcd. for $C_{27}H_{43}N_6O$: 467.3498; found: 467.3489.

**4-Chloro-6-methoxy-2-(piperidin-1-yl)-7-[3-(piperidin-1-yl)propoxy]quinazoline (8):** mp.: 138 °C. $R_f$ = 0.35 (15% MeOH in $CH_2Cl_2$ IR (film): ṽ = 2931, 1626, 1585, 1495, 1238, 754 $cm^{-1}$. $^1$H NMR (500 MHz, $CD_3OD$): $\delta$ = 1.59-1.70 (m, 6 H, $CH_2CH_2CH_2NCH_2CH_2CH_2O$, $CH_2CH_2NC$), 1.70-1.77 (m, 2 H, $CH_2CH_2CH_2NC$), 1.77-1.89 (m, 4 H, $CH_2CH_2NCH_2CH_2CH_2O$), 2.15-2.37 (m, 2 H, $CH_2CH_2O$), 2.87-3.17 (m, 6 H, $CH_2NCH_2CH_2CH_2O$), 3.81-3.88 (m, 4 H, $CH_2NC$), 3.95 (s, 3 H, $CH_3O$), 4.25 (t, *J* = 5.8 Hz, 2 H, $CH_2O$), 7.00 (s, 1 H, $CHCOCH_2$), 7.26 (s, 1 H, $CHCOCH_3$). $^{13}$C NMR (126 MHz, $CD_3OD$): $\delta$ = 23.72 ($CH_2CH_2CH_2NCH_2CH_2CH_2O$), 25.29 ($CH_2CH_2NCH_2CH_2CH_2O$), 25.74 ($CH_2CH_2O$)*, 25.92 ($CH_2CH_2CH_2NC$)*, 26.88 ($CH_2CH_2NC$), 46.31 ($CH_2NC$), 55.01 ($CH_2NCH_2CH_2CH_2O$), 56.54 ($CH_3O$), 56.73 ($CH_2CH_2CH_2O$), 68.09 ($CH_2O$), 104.82 ($CHCOCH_3$), 106.56 ($CHCOCH_2$), 113.18 (CCCl), 148.82 ($COCH_3$), 152.67 (CCCCl), 157.42 ($COCH_2$), 158.93 ($CNCH_2$), 160.90 (CCl). HRMS-ESI m/z $[M+H]^+$ calcd. for $C_{22}H_{32}ClN_4O_2$: 419.224, found: 419.2206.

**4-Chloro-6-methoxy-7-[3-(piperidin-1-yl)propoxy]quinazoline (9)** [6, 7]**:** mp.: 106 °C. $R_f$ = 0.21 [5% 4 M $NH_3$ (in MeOH) in $CH_2Cl_2$]. IR (film): ṽ = 2933, 2358, 1500, 1232, 1020 $cm^{-1}$. $^1$H NMR (500 MHz, DMSO-$d_6$) $\delta$ = 1.31-1.43 (m, 2 H, $CH_2CH_2CH_2NCH_2CH_2CH_2O$), 1.45-1.54 (m, 4 H, $CH_2CH_2NCH_2CH_2CH_2O$), 1.95 (p, *J* = 6.7 Hz, 2 H, $CH_2CH_2O$), 2.26-2.38 (m, 4 H, $CH_2NCH_2CH_2CH_2O$), 2.38-2.43 (m, 2 H, $CH_2CH_2CH_2O$), 4.00 (s, 3 H, $CH_3O$), 4.25 (t, *J* = 6.4 Hz, 2 H, $CH_2O$), 7.38 (s, 1 H, CHCN), 7.44 (s, 1 H, CHCCN), 8.86 (s, 1 H, CHN). $^{13}$C NMR (126 MHz, DMSO-$d_6$) $\delta$ = 24.13 ($CH_2CH_2CH_2NCH_2CH_2CH_2O$), 25.60 ($CH_2CH_2NCH_2CH_2CH_2O$), 25.92 ($CH_2CH_2O$), 54.11 ($CH_2NCH_2CH_2CH_2O$), 54.93 ($CH_2CH_2CH_2O$), 56.21 ($CH_3O$), 67.65 ($CH_2O$), 102.30 (CHCCN), 107.32 (CHCN), 118.48 (CCCl), 148.60 (CCCCl), 151.49 ($CH_3OC$), 152.19 (CHN), 156.11 ($CH_2OC$), 157.84 (CCl). HRMS-ESI m/z $[M+H]^+$ calcd. for $C_{17}H_{23}ClN_3O_2$: 336.1479, found: 336.1476. The analytical data agree with those previously reported in the literature [6, 7].

**7-Fluoro-*N*-(1-propan-2-ylpiperidin-4-yl)-6-methoxyquinazolin-4-amine (10):** mp.: 233 °C (decomposition). $R_f$ = 0.48 [10% 3 M $NH_3$ (in MeOH) in $CH_2Cl_2$]. IR (film): ṽ = 2968, 1593, 1537, 1045 $cm^{-1}$. $^1$H NMR (500 MHz, $CD_2Cl_2$): $\delta$ = 1.04 (d, *J* = 6.6 Hz, 6 H, $CH_3CH$), 1.52-1.64 (m, 2 H, $CH_2CHNH$), 2.11-2.20 (m, 2 H, $CH_2CHNH$), 2.31-2.42 (m, 2 H, $CH_2CH_2CHNH$), 2.71-2.84 (m, 1 H, $CH_3CH$), 2.83-2.96 (m, 2 H, $CH_2CH_2CHNH$), 4.00 (s, 3 H, $CH_3O$), 4.15-4.31 (m, 1 H, CHNH), 5.43 (d, *J* = 7.6 Hz, 1 H, NH), 7.02 (d, *J* = 8.6 Hz, 1 H, $CHCOCH_3$), 7.44 (d, *J* = 12.0 Hz, 1 H, CHCF), 8.48 (s, 1 H, CHNCNH). $^{13}$C NMR (126 MHz, $CD_2Cl_2$): $\delta$ = 18.43 ($CH_3CH$), 33.01 ($CH_2CHNH$), 48.00 ($CH_2CH_2CHNH$), 49.23 (CHNH), 54.86 ($CH_3CH$), 56.96 ($CH_3O$), 102.29 (d, *J* = 3.5 Hz, $CHCOCH_3$), 111.90 (CCNH), 114.04 (d, *J* = 17.6 Hz, CHCF), 146.26 (d, *J* = 12.0 Hz, CCHCF), 147.82 (d, *J* = 13.0 Hz, $COCH_3$), 154.79 (CHNCNH), 156.64 (d, *J* = 255.1 Hz, CF), 158.12 (CNH). HRMS-ESI m/z $[M+H]^+$ calcd. for $C_{17}H_{24}FN_4O$: 319.1934, found: 319.1929.

* Due to signal overlap in 2D-NMR, signals could not be unambiguously assigned.
